# FXR and BET signaling orchestrate to protect β cells

**DOI:** 10.64898/2026.04.10.716420

**Authors:** Fritz Cayabyab, Jalan Tipirneni, David Chen, Jinhyuk Choi, Yu Hamba, Nhi Pham, Clarissa Tacto, Jie Wu, Liu Wang, Yeva Mirzakhanyan, Paul David Gershon, Harvey Perez, Naoki Harada, Kiyoka Kim, Ameen Shaheen, Sungsoon Fang, Eli Ipp, Lin-Feng Chen, Zong Wei, Eiji Yoshihara

**Author notes:** Correspondence Keyword; FXR, BRD4, *db/db*, β cells, HILOs.

## Abstract

In both type 1 and type 2 diabetes (T1D and T2D), insulin-producing β cells undergo progressive dysfunction due to inflammation, leading to impaired glucose responsiveness, dedifferentiation, and cell loss. While bile acid (BA) dysregulation under diabetic conditions is known to influence metabolic and inflammatory pathways, its mechanistic role in β cell regulation remains incompletely defined^1–3^. Here we show that bile acid sensor Farnesoid X receptor (FXR) and Bromodomain and Extra-Terminal motif (BET) signaling cooperatively regulates β cell inflammatory response and β cell identity. We identified the physiological protein-protein interaction between FXR and the bromodomain-containing protein 4 (BRD4) as a regulatory axis that protects against β cell dysfunction. We show that FXR activation by Fexaramine (Fex) together with BRD4 inhibition by JQ1 synergistically suppressed IL-1β-induced inflammation while also improving β cell identity and insulin secretion in both *db/db* model and high-fat diet (HFD) plus multi low-dose streptozotocin (MLD-STZ) model of diabetes. Importantly, this cooperative effect is abolished in β cell–specific FXR knockout (βFXRKO) mice, establishing that FXR is required for the functional synergy between these pathways *in vivo*. Mechanistically, structure-guided modeling and mutational analyses identified a direct interaction between FXR and the BD2 domain of BRD4, depending on specific lysine acetylation sites. Additionally, inhibition of the BD2 domain of BET combined with FXR activation markedly improved β cell survival in human T1D and T2D models established from human pluripotent stem cell (hPSC)-derived islet-like organoids (HILOs). Collectively, these findings establish a BA–bromodomain axis as a transcriptional interface linking metabolic signaling and chromatin regulation, and highlight FXR–BET targeting as a promising strategy to counter progressive β cell failure in diabetes.

## Introduction

Deterioration of β cell mass and function is a primary cause of both type 1 diabetes (T1D) and type 2 diabetes (T2D)^4^. Although exogenous insulin therapy remains the cornerstone of treatment, it neither halts disease progression nor restores β cell health. Therapeutic strategies aimed at preserving or restoring β cell function are therefore of high priority. Emerging evidence has implicated bile acids (BAs) as modulators of β cell inflammation and dysfunction, with altered BA composition observed in both T1D^5–9^ and T2D^1–3,8,10^. Yet, the mechanisms by which BAs regulate β cell fate remain incompletely understood.

The nuclear receptor FXR is a key BA sensor that is widely expressed in various cell types. It regulates lipid metabolism and suppresses inflammation in multiple tissues, including the liver and intestine^11–24^. FXR is also expressed in pancreatic islets, where it influences insulin secretion and β cell gene expression^21,25–27^. The BA pool comprises both FXR agonists and antagonists, and synthetic ligands such as Fexaramine (Fex) offer a strategy to selectively activate FXR^28^. Specific BAs differentially regulate FXR activity with species such as chenodeoxycholic acid (CDCA) and cholic acid (CA) acting as potent agonists^29^, whereas others, including muricholic acids particularly tauro-conjugated form (βTMCA)^24,30,31^, suggesting that BA composition shapes FXR signaling and cellular function. Certain hydrophilic BAs, such as tauroursodeoxycholic acid (TUDCA), primarily act as chemical chaperones that alleviate endoplasmic reticulum (ER) stress, whereas deoxycholic acid (DCA) exhibit context-dependent effects on FXR signaling^24,32^. Pharmacological activation of FXR using Fex has been demonstrated to ameliorate tissue inflammation and restore metabolic homeostasis in both intestine-specific via oral gavage^24,33^ and systemic administration by intraperitoneal (i.p.) injection^34^. However, the precise role of FXR signaling in β cells during diabetes progression, and how it interacts with other regulatory networks, is poorly defined.

The bromodomain-containing protein (BRD) family consists of well-known chromatin reader proteins that recognize acetyl (Ac) lysine of histones and non-histone proteins. This interaction plays a pivotal role in regulating gene expression, including inflammatory responses in many cell types, making it a favorable target for therapeutic development^35–38^. Bromodomain and Extra-Terminal motif (BET) inhibitors, such as the well-characterized JQ1, selectively inhibit BRD2, BRD3, and BRD4^39^, showing potential in reducing cancer proliferation and inflammation^35–38^. Notably, BRD4 is known as an activator of NF-κB-mediated inflammatory gene expression in various cells, including epithelial cells and macrophages^36,40,41^. It also influences metabolic processes such as liver cholesterol metabolism^42^, adipose lipolysis through tissue resident macrophage^43^ and insulin secretion in β cells^44^. Our protein purification experiments identified an interaction between FXR and BRD family proteins. Given the established roles of FXR and BRD4 in the reciprocal regulation of inflammation and maintenance of key β cell identity transcription factors, we hypothesized that the BA sensor FXR and epigenetic readers BRDs cooperatively regulate inflammation and cellular identity in β cells during the pathogenesis of diabetes. To test this hypothesis, we employed a genetically engineered human β cell line (EndoC-βH1 cells)^45^, a severe T2D mouse model (*db/db* mice and HFD + MLD-STZ mice), and the HILOs^46^. β cell plasticity is dysregulated by inflammatory cytokines such as IL-1β, IFNγ and IFNβ during the progression of human T2D^47^, which is partially mimicked in the leptin receptor-deficient *db/db* mice. Inflammatory cytokines trigger β cell dedifferentiation, characterized by a reduction in insulin gene and urocortin 3 (UCN3) expression^48,49^, followed by β cell endoplasmic reticulum (ER) stress and apoptosis^47^. In the context of human T2D, accumulation of thioredoxin-interacting protein (TXNIP) and human islet amyloid polypeptide (IAPP) in β cells leads to the cytokine/ER-stress-mediated β cell dysfunction^50^. HILOs provide a unique, scalable model for human islet generation and disease modeling^46^. To capture distinct pathological features of diabetes, we established complementary *in vitro* models of T2D and T1D using HILOs. We induced human TXNIP and human IAPP overexpression in HILO to recapitulate immune-metabolic stress-driven β cell dysfunction as an *in vitro* human T2D model. In parallel, we co-cultured MHC-matched HILOs and human T1D-derived PBMC (T1DPBMC) as an *in vitro* human T1D model.

This work redefines how nuclear receptor and chromatin pathways converge in β cells, establishing a bile acid–epigenetic axis as a central regulator of diabetes progression and a promising therapeutic target for preserving β cell function.

## Results

### Dysregulation of bile acid signaling is implicated in the β cell dysfunction in diabetes

We previously observed that high-fat diet (HFD) alters the BA production and composition in mice with or without genetic mutation^24,33^. Building on these findings, we investigated whether BA alternations contribute to the pathogenesis of diabetes beyond the context of HFD. To this end, we performed targeted metabolomic analyses of serum BAs across three diabetic mouse models, HFD + MLD-STZ (T2D), leptin receptor-deficient *db/db* (T2D) and non-obese spontaneous diabetic NOD (T1D). Serum BA profiling revealed that both total BA levels and BA composition were altered across all three models during the progression of hyperglycemia (Supplementary Fig. 1a-c, Fig. 1a). In HFD + MLD-STZ mice, CDCA, DCA, lithocholic acid (LCA), βMCA, ω-muricholic acid (ωMCA), taurodeoxycholic acid (TDCA) and ursodeoxycholic acid (UDCA) were significantly increased compared to age- and sex- matched controls. Similarly, in male *db/db* mice, multiple BAs including DCA, hyodeoxycholic acid (HDCA), βMCA, ωMCA, taurocholic acid (TCA), taurochenodeoxycholic acid (TCDCA), tauroursodeoxycholic acid (TUDCA) and UDCA were increased compared to BKS or C57BL/6J controls (Fig.1b). In female NOD mice at 19 weeks of age, TCA and α+βTMCA levels were significantly increased in diabetic compared to non-diabetic mice, whereas TDCA and TUDCA showed a non-significant trend toward increase (Fig.1c). In addition, longitudinal analyses revealed that DCA, ωMCA, and TDCA levels were significantly increased at 23 weeks compared to 8 weeks in female NOD mice (Supplementary Fig. 1d, e). Together, these results indicate that BA dysregulation is a common feature across both T1D and T2D models. FXR is a key BA receptor that is regulated by BA composition^11,12,15–20,22–24^ and is downregulated in islets of T2D models including *db/db* mice^51^. Although β cells play a critical role in glucose homeostasis in these models, the role of FXR signaling in β cells, particularly under the stress of disease progression remains unclear. Consistent with the previous reports^21,25–27,52–54^, we confirmed that FXR is expressed in human islets and EndoC-βH1 cells and is upregulated during human β cell differentiation in HILOs^46^ (Supplementary Fig. 2a-d). Treatment with FXR ligands (Fex or GW4046) increased FXR responsive cis-element (FXRE) reporter activity in EndoC-βH1 cells and HEK293LTV cells, confirming functional FXR signaling (Supplementary Fig. 2e). In contrast, Fex did not stimulate TGR5-dependent secreted alkaline phosphatase (SEAP) reporter activity, indicating specific for FXR signaling (Supplementary Fig. 2f). We next examined how individual BAs modulate FXR activity. Acute exposure (24 hours) to glycohyocholic acid (GHCA) and CDCA increased FXRE activity, whereas chronic exposure (72 hours) to hyocholic acid (HCA), βMCA, hyodeoxycholic acid (HDCA), taurohyodeoxycholic acid (THDCA), and TCDCA suppressed FXRE activity (Supplementary Fig. 2g). Functionally, chronic BA-exposure including TUDCA, βMCA, CA, TCA, CDCA, DCA and HDCA increased sensitivity to cytokine (IL-1β + IFNγ)-induced β cell apoptosis, as measured by caspase 3/7 activation in EndoC-βH1 cells (Fig. 1d, e). In contrast, Fex treatment ameliorated cytokine- and BA-induced apoptosis (Fig. 1f), while BAs alone had minimal effects in the absence of cytokine stimulation (Supplementary Fig. 2h). These findings suggest that chronic BA exposure can modulate β cell inflammatory responses through FXR signaling.

**Fig. 1.**
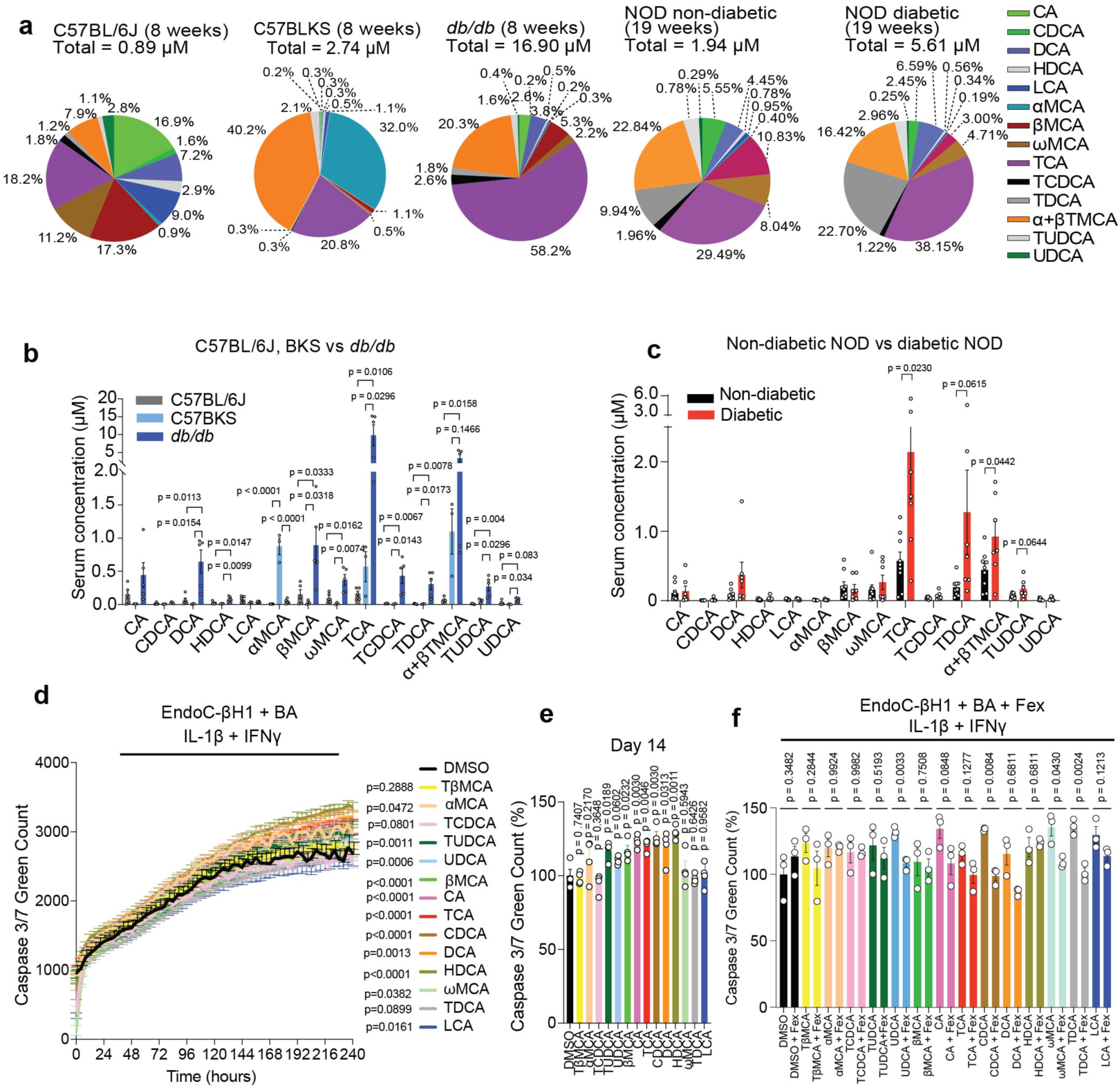
Dysregulation of bile acid signaling in T1D and T2D model mice. **a**, Comparison of serum BA profile as total and average percent composition in C57BL/6J mice (8 weeks, male, n=5), C57BLKS mice (8 weeks, male, n = 3) and *db/db* mice (8 weeks, male, n = 5), non-diabetic NOD mice (19 weeks, female, n = 9), diabetic NOD mice (19 weeks, female, n = 7). **b,** Comparison of serum individual BA amount in C57BL/6J (non-diabetic control, n = 5), C57BLKS (non-diabetic control, n = 3) and *db/db* (diabetic, n = 5). c, Comparison of serum individual BA amount in non-diabetic (n = 9) and diabetic (n = 7) female NOD mice at 19 weeks of age. **d,** Caspase 3/7 green count in EndoC-βH1 cells for 240 hours treatment with different BAs in the presence of IL-1β (10 ng/ml) and IFNγ (10 ng/ml). n = 4/each. **e**, Caspase 3/7 green count in EndoC-βH1 cells at 14 days treatment with different BAs in the presence of IL-1β (10 ng/ml) and IFNγ (10 ng/ml). n = 4/each. **f**, Caspase 3/7 green count in EndoC-βH1 cells at 240 hours treatment after addition of Fex (10 μμ) in the presence of BAs (10 μM each), IL-1β (10 ng/ml) and IFNγ (10 ng/ml) (n = 3/each). Statistical analyses were performed using one-way ANOVA with Tukey’s multiple-comparison test per bile acid (BA) condition for (b, e), unpaired two-tailed Student’s t-test per BA condition for (c, f), and two-way ANOVA for (d). Error bars represent mean ± SEM.

### Physical interaction between FXR and BRD4

To elucidate the molecular mechanism underlying FXR-mediated protection, we investigated whether FXR physically interacts with chromatin regulatory proteins. Proteomic pull-down analysis using V5-tagged FXR as bait identified multiple bromodomain-containing proteins, including BRD2, BRD3, and BRD4, as FXR-interacting partners in HEK293LTV cells (Fig. 2a). Given the established role of BRD4 in inflammatory transcriptional regulation, we focused on this interaction. As recognized epigenetic regulators, BETs modulate chromatin modifications and inflammatory responses, playing pivotal roles across various pathophysiological conditions ^37,38,42,55,56^. Nuclear receptors (NRs) are known to interact with BET family proteins, which influences their transcriptional activity during the inflammatory responses^54,57^. Co-immunoprecipitation (IP) analyses across multiple systems, including HEK293LTV, EndoC-βH1, and INS-1 cells, consistently demonstrated a robust interaction between FXR and BRD4 (Fig. 2b–e, Supplementary Fig.3a, b). This interaction was significantly reduced upon treatment with either the FXR agonist Fex or the BET inhibitor JQ1, indicating that FXR–BRD4 binding is dynamically regulated by ligand signaling. A previous study indicates that K158 and K218 lysine sites are functional acetylation sites of FXR^58^. Because bromodomains bind acetylated lysines, we hypothesized that the acetylation of FXR, specifically K158 and K218 is required for the interaction with BRD4. Supporting this notion, structure prediction using AlphaFold Chimera-based modeling identified a direct interaction between FXR containing K158 and K218 region and the BD2 domain of BRD4, mediated through lysine acetylation sites (Fig. 2f). Indeed, acetylation (Ac) of FXR protein is synergistically decreased by Fex + JQ1 treatment in EndoC-βH1 cells (Fig. 2g). To confirm the involvement of K158Ac and K218Ac in the FXR-BRD4 interaction, we mutated K158 and/or K218 to arginine (K158R and/or K218R). As predicted, we found that these double mutations of K158 and K218 significantly decreased the lysine acetylation of FXR and the interaction with BRD4 in EndoC-βH1 and INS-1 cells (Fig. 2h and Supplementary Fig. 3a). Although p300 and RXR are significant co-activators of FXR, overexpressing those cofactors did not rescue the interaction between FXR and BRD4 interaction (Supplementary Fig. 3b), suggesting that FXR’s lysine-acetylation sites may be critical for direct binding to BRD4. To assess whether the disruption of FXR–BRD4 interaction was due to non-specific lysine substitution, we generated additional FXR mutants. Mutations at K335 and K420, located within the hinge–LBD region involved in co-regulator recruitment, did not affect BRD4 binding. These results indicate that the FXR–BRD4 interaction depends on specific lysine residues rather than general lysine modification (Supplementary Fig. 3c). Collectively, our results indicate a ligand-dependent mechanism for FXR that involves the recruitment and release of lysine acetylation reader BRD4 in β cells.

**Fig. 2.**
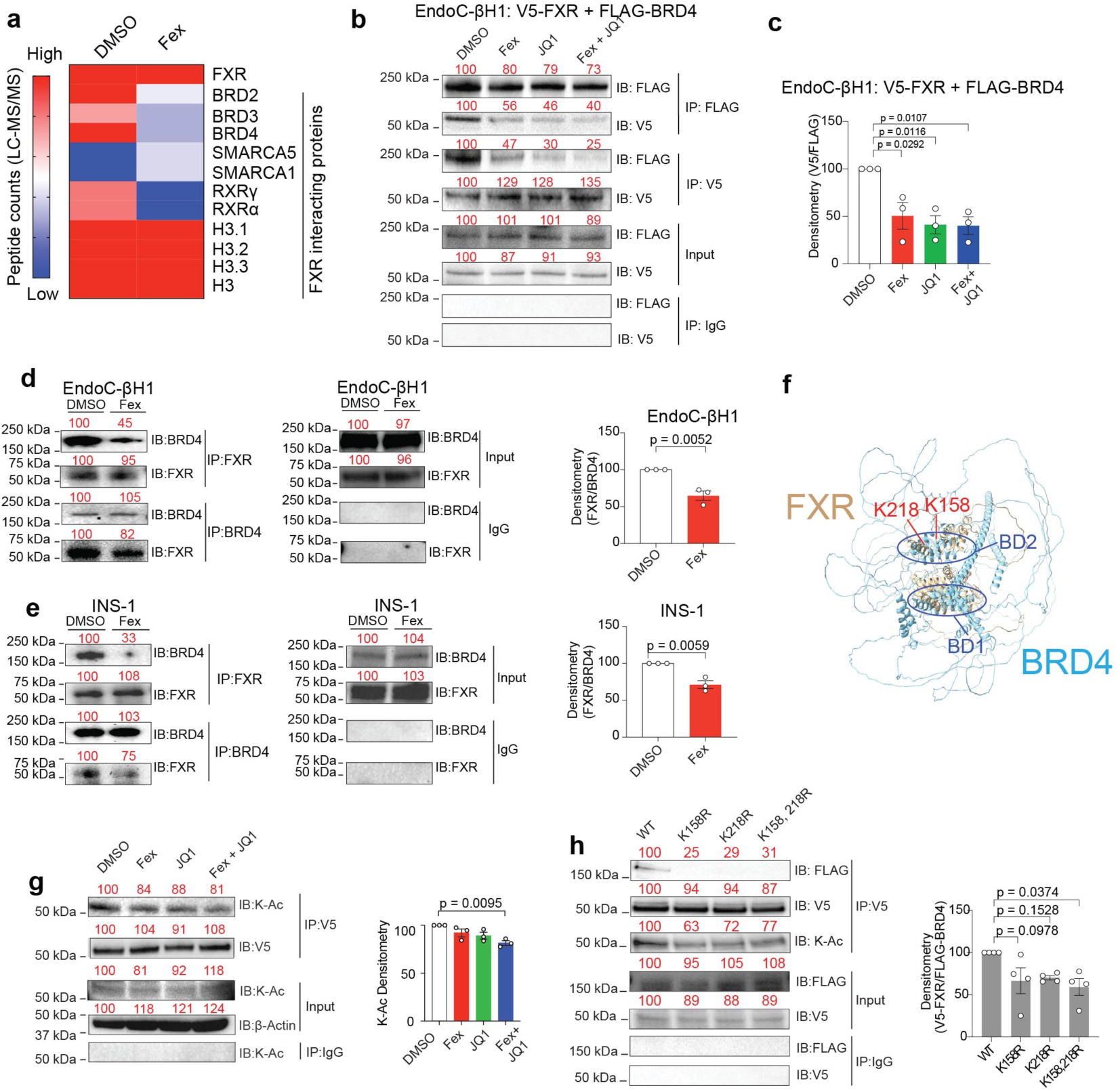
Physical interaction between FXR and BRD4 through FXR acetylated Lysine. a, Heatmap analyses of peptide counts (LC-MS/MS) proteins after antibody-based protein purification of FXR in HEK293LTV cells after treatment with DMSO or Fex (10 μM) for 24 hours. **b,** Co-immunoprecipitation (co-IP) of V5-FXR and FLAG-BRD4 after V5 pull-down (IP:V5) or FLAG pull-down (IP:FLAG) in EndoC-βH1 cells treated with DMSO, Fex, JQ1 and Fex + JQ1. Indicated immunoblot (IB) was performed. IgG pulldown control and input also shown. Red numbers above the blots indicate densitometry values presented as percent relative to DMSO which is set as 100. **c,** Densitometric analyses of V5-FXR/FLAG-BRD4 ratios in EndoC-βH1 cells treated with DMSO, Fex, JQ1 or Fex+JQ1 from three independent co-IP experiments. Densitometric analyses of FXR/BRD4 ratio were shown as %. (**d, e**) Co-IP of FXR (IP:FXR) with BRD4 or BRD4 (IP:BRD4) with FXR in EndoC-βH1 cells (d) or INS-1 cells (e). Red numbers above the blots indicate densitometry values presented as percent relative to DMSO which is set as 100. Bar graph on the right represents densitometric analyses of FXR/BRD4 ratios in EndoC-βH1 cells treated with DMSO or Fex from three independent co-IP experiments. **f**, Protein structure modeled by Alphafold Chimera X. Possible interaction between lysine acetylation site in FXR (K158, K218) and BD2 domain in BRD4 are highlighted. **g**, Immunoblot (IB) of acetylated lysine (K-Ac) or V5 in FXR pull-down (IP:V5) protein complexes from EndoC-βH1 cells. Red numbers above the blots indicate densitometry values presented as percent relative to DMSO which is set as 100. The bar graph represents densitometric analyses of K-Ac/V5-FXR ratios from three independent experiments. **h**, Immunoblot (IB) of FLAG, V5 or Acetylated lysine (K-Ac) in V5-FXR (WT), V5-FXR (K158R), V5-FXR (K218R), V5-FXR (K158R/K218R) pull-down (IP:V5) protein complexes from EndoC-βH1 cells. Red numbers above the blots indicate densitometry values presented as percent relative to FXR-WT which is set as 100. Statistical analyses were performed using one-way ANOVA with Tukey’s multiple-comparison test for (c, g, h) and unpaired two-tailed Student’s t-test for (d, e). Error bars represent mean ± SEM.

### FXR agonist and BET inhibitor synergistically regulate β cell transcriptome and insulin secretion under inflammatory stress

To investigate the physiological role of FXR and BET signaling in β cells, we first examined whether BRD4 modulates the anti-inflammatory function of FXR. To determine the role of FXR in islets, we isolated islets from Fxr flox/flox mice^59^ and deleted the *Fxr ex vivo* using adenovirus-mediated Cre recombination (Ad-FxrKO), with GFP-expressing virus as control (Ad-Control) (Supplementary Fig. 4a). IL-1β and/or IFNγ are often used to induce inflammatory responses in islets. Consistent with a protective role of FXR, Fex partially rescued the IL-1β + IFNγ-induced impairment of glucose-stimulated insulin secretion (GSIS) in ad-Control islets, whereas Fex failed to rescue the IL-1β + IFNγ-mediated defect of GSIS in ad-FxrKO islets (Fig. 3a). To determine the role of Brd4 in islets, we isolated islets from Brd4 flox/flox mice^43^ and performed *Brd4* gene knockout by adenovirus-mediated Cre expression (Ad-Brd4KO) or control GFP expression (ad-Control) *ex vivo* (Supplementary Fig. 4b). We found that Ad-Brd4KO islets ameliorated cytokine-induced GSIS impairment, suggesting that FXR and BRD4 exert opposing roles in regulating β cell function under inflammatory stress (Fig. 3b). We next evaluated insulin secretion across multiple systems, including primary mouse and human isolated islets, as well as EndoC-βH1 cells expressing proinsulin-linked Gaussia luciferase reporter (Proinsulin-NanoLuc)^60^. In both mouse and human islets, combined Fex and JQ1 treatment synergistically restored GSIS impaired by IL-1β + IFNγ (Fig. 3c, d). In EndoC-βH1 cells, cytokine-induced dysfunction of insulin secretion was more robustly triggered by IL-1β + IFNβ than IL-1β + IFNγ (Supplementary Fig. 4c). Given the limited glucose responsiveness of EndoC-βH1 cells, insulin secretion was assessed under depolarizing conditions (KCl), where Fex + JQ1 treatment similarly restored cytokine-impaired secretion (Fig. 3e, Supplementary Fig. 4d). To define the transcriptional programs regulated by FXR and BET signaling, we performed bulk RNA-seq in EndoC-βH1 cells treated with Fex, JQ1, or both, in the presence or absence of IL-1β. IL-1β stimulation upregulated 4,359 genes, among which 924 and 1,920 genes were suppressed by Fex and JQ1, respectively (Fig. 3f). Notably, 684 genes were commonly downregulated, and 186 of these (including *HLA-A*, *CXCL2*, *TNFRSF11B*, and *CASP4*) were synergistically suppressed by combined treatment. qPCR validation confirmed that this synergistic repression observed in RNA-seq under IL-1β stimulation is preserved under combined IL-1β + IFNγ treatment (Supplementary Fig. 4e). Gene ontology (GO) analysis and Gene Set Enrichment Analysis (GSEA) analyses indicated enrichment of pathways related to apoptosis, immune response, and ER stress (Fig. 3g, Supplementary Table 1). Consistent with these results, we found characteristic binding motifs for the known inflammatory response transcription factors STAT3, JUN, VDR, and NF-κB in the regulatory regions of the genes downregulated by Fex + JQ1 in EndoC-βH1 cells (Fig. 3h, Supplementary Table 2). FXR signaling is known to suppress the inflammatory response through NF-κB signaling^61^. Next, we performed a firefly luciferase assay in which luciferase was expressed under the control of an NF-κB promoter (NF-κB-Luc) to evaluate FXR-mediated inflammation suppression in HEK293LTV and EndoC-βH1. In both HEK293LTV and EndoC-βH1 cells, we observed that Fex or JQ1 individually suppressed NF-κB-Luc activity in the presence of IL-1β, while in combination, Fex and JQ1 exhibited a synergistic suppression of NF-κB-Luc activity (Fig. 3i). Notably, JQ1 did not affect FXRE reporter activation induced by Fex (Supplementary Fig. 4f), whereas siRNA-mediated knockdown of BET proteins further reduced NF-κB activity (Supplementary Fig. 4g, h), suggesting that BET proteins act redundantly to promote inflammatory transcription.

**Fig. 3.**
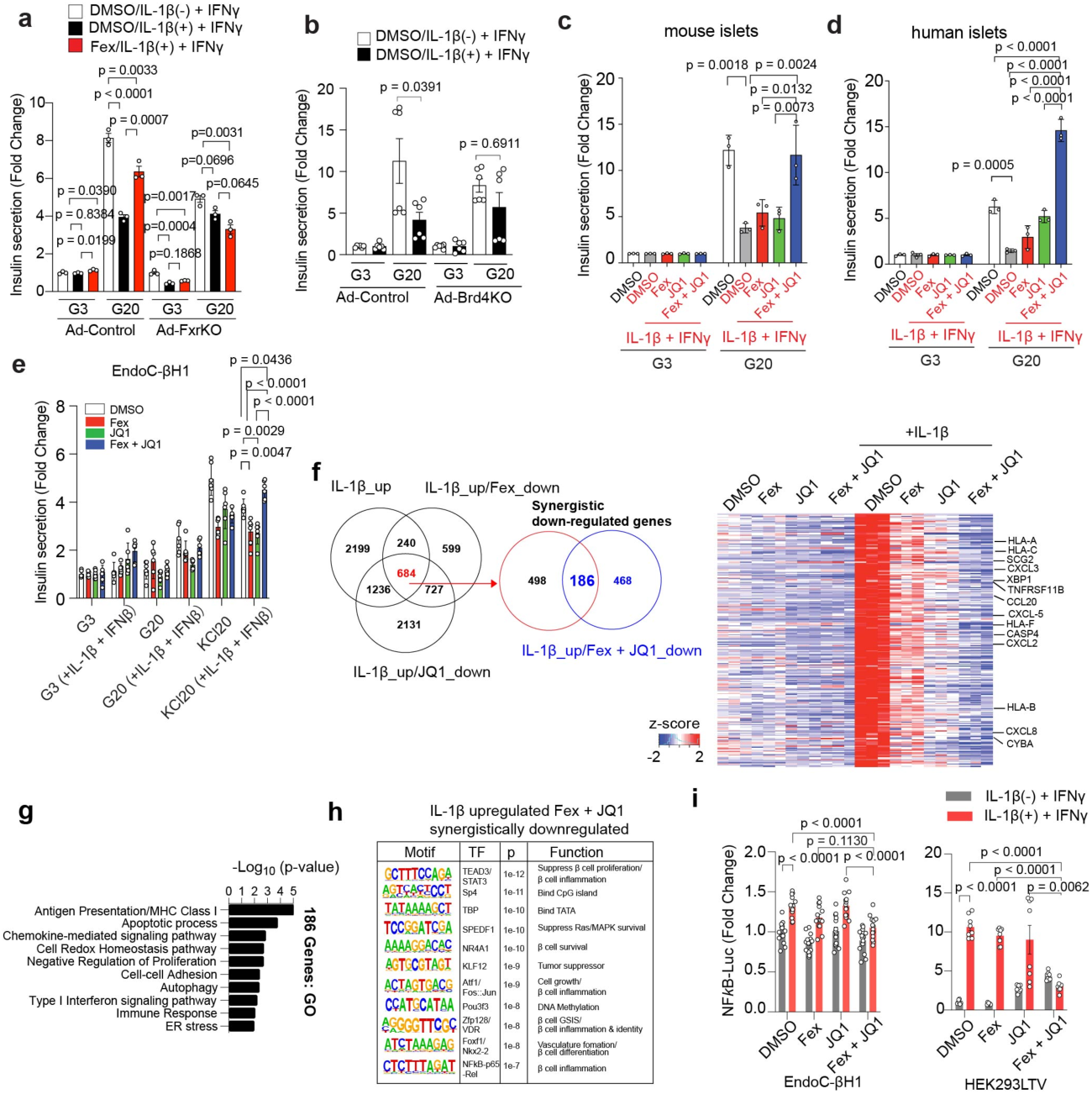
FXR agonist and BET inhibitor synergistically regulate β cell transcriptome and insulin secretion under the inflammatory stress. **a**, GSIS assay of isolated islets from Fxr flox/flox mice. Adenovirus-mediated GFP (Ad-Control) or Cre-GFP (Ad-FxrKO) expression was performed 96 hours prior to the GSIS assay. Fex (10 μM) were pretreated 72 hours prior to the GSIS and IL-1β (10 ng/ml) + IFNγ (10 ng/ml) were pretreated 48 hours prior to the GSIS assay. G3 = glucose 3 mM, G20 = glucose 20 mM. Fold change over control (Ad-Control G3) is shown. 10 islets per condition, n = 3. **b**, GSIS assay of isolated islets from Brd4 flox/flox mice. Adenovirus-mediated GFP (Ad-Control) or Cre-GFP (Ad-Brd4KO) expression was performed 72 hours prior to the GSIS assay. Mouse islets were pretreated with IL-1β (10 ng/ml) + IFNγ (10 ng/ml) 48 hours prior to the GSIS assay. G3 = glucose 3 mM, G20 = glucose 20 mM. Fold change over control (Ad-Control G3) is shown. 10 islets per condition, n = 6. **c**, GSIS assay of isolated islets from C57BL/6J mice. Mouse islets were pretreated with Fex (10 μM), JQ1 (500 nM), or Fex (10 μM) +JQ1 (500 nM) for 72 hours prior to the GSIS assay. In addition, the islets were pretreated with IL-1β (10 ng/ml) + IFNγ (10 ng/ml) for 48 hours prior to the GSIS assay. 10 islets per condition, n = 3. **d**, GSIS assay of primary human islets. Human islets were pretreated with DMSO, Fex (10 μM), JQ1 (500 nM), or Fex (10 μM) +JQ1 (500 nM) for 72 hours prior the GSIS assay. In addition, the islets were pretreated with IL-1β (10 ng/ml) + IFNγ (10 ng/ml) for 48 hours prior to the GSIS assay.10 islets per condition, n = 3. **e**, GSIS assay in EndoC-βH1 after 72 hours pretreatment with DMSO, Fex (10 μM), JQ1 (500 nM), or Fex (10 μM) +JQ1 (500 nM) and 48 hours treatment with IL-1β (10 ng/ml) and IFNβ (10 ng/ml). 100,000 cells/well for n = 6/each. **f**, Venn diagram of differentially expressed genes in EndoC-βH1 cells pretreated with DMSO, Fex (10 μM), or JQ1 (500 nM) for 72 hours and with IL-1β (10 ng/ml) for 48 hours. n = 3/each. Heatmap of z-score for the 186 synergistic genes regulated by Fex and JQ1 in EndoC-βH1 cell. RNA-seq analyses were performed after treatment with DMSO, Fex (10 mM), JQ1 (500 nM), or Fex (10 μM) +JQ1 (500 nM) for 72 hours and with IL-1β (10 ng/ml) for 48 hours. n = 3/each. **g**, Gene ontology of the 186 synergistic genes. n=3. **h**, Promoter motif analysis of the 186 synergistic genes. n = 3. **i**, NFκB-luciferase reporter assay in EndoC- βH1 and HEK293LTV cells after 48 hours treatment with DMSO, Fex (10 μM), JQ1 (500 nM), or Fex (10 μM) + JQ1 (500 nM) and 24 hours treatment with IL-1β (10 ng/ml). Statistical analyses were performed using unpaired two-tailed Student’s t-test for (b) and one-way ANOVA with Tukey’s multiple-comparison test for (a, c–e, i). Error bars represent mean ± SEM.

Given the importance of chromatin accessibility in inflammatory gene regulation, we next performed ATAC-seq under IL-1β stimulation with DMSO, Fex, JQ1, or Fex + JQ1 treatment (Supplementary Fig. 5a). While global chromatin accessibility was not significantly altered between groups (Supplementary Fig. 5b), IL-1β stimulation significantly increased (> 2-fold) in chromatin accessibility at 4,437 gene related loci and Fex and JQ1 commonly decreased (< 2-fold) 1,048 genes (∼23.6%) at same loci (Supplementary Fig. 5c). In contrast, IL-1β stimulation significantly decreased (< 2-fold) in chromatin accessibility at 2,450 gene related loci and Fex and JQ1 co-treatment restored accessibility (> 2-fold) at 121 genes (∼4.9%) at same loci (Supplementary Fig. 5c). GO analyses for selectively responsive to Fex + JQ1 (compared to Fex or JQ1 alone under IL-1β stimulation) revealed upregulation of β cell function–related pathways, such as cAMP signaling and sodium ion transport, and downregulation of inflammatory pathways including NF-κB activity and leukocyte activation (Supplementary Fig. 5d, e). Notably, chromatin accessibility at *STAT3*, *STAT5B*, and *NF-κB* loci was synergistically reduced by Fex + JQ1 treatment (Supplementary Fig. 5f). These results implicate a direct role of FXR and BRD4 in counteracting the effect of IL-1β-mediated inflammatory response. To further examine chromatin-level regulation, we performed BRD4 ChIP-seq under IL-1β stimulation with or without Fex. Fex treatment increased BRD4 binding peaks from 4,868 to 6,629, accompanied by reduced enrichment of inflammatory transcription factor motifs such as SMAD3 (Supplementary Fig. 5g–i). Collectively, these results support a model in which FXR activation and BET inhibition cooperatively reprogram β cell chromatin landscapes, suppress inflammatory gene expression, and preserve β cell identity under cytokine stress.

### FXR agonist and BET inhibitor ameliorate hyperglycemia in *db/db* mice

Improving β cell function in progressive T2D remains a major therapeutic challenge. While FXR activation and BET inhibition have each independently been shown to improve glucose homeostasis in HFD-induced obese models^33,43^, it remains unclear whether combined therapy targeting FXR and BET proteins can ameliorate advanced β cell failure. To address this, we utilized *db/db* mice on a BKS background, a model characterized by pancreatic β cell loss, hypoinsulinemia, and hyperglycemia. Fex (50 mg/kg) and/or JQ1 (25 mg/kg) were administered via intraperitoneal (i.p.) injection three times weekly starting at 5 weeks of age. Single-dose Fex administration activated target genes across multiple tissues such as *Mafa*, *Ucn3* and *G6pc2* in islets, *Dio2*, *Esrrg*, *Ppargc1a*, *Ppargc1b*, *Slc2a4* and *Ucp1* in brown adipose tissue (BAT), *Nr1h4* (*Fxr*), *Shp*, *Cyp7a1* and *Cyp8b1* in liver and *Fabp6*, *Fgf15*, *Osta* and *Bsep* in Ileum of intestine in C57BL/6J mice (Supplementary Fig. 6a, b). These results indicate that i.p. injection of Fex can circulate peripheral metabolic tissues. While single administration with Fex or JQ1 did not significantly alter fasting glucose levels, combined treatment significantly reduced fasting glucose at 12 and 13 weeks (Fig. 4a, Supplementary Fig. 6c). Similarly, fed *ad lib* glucose levels were decreased and serum insulin levels were increased in the Fex + JQ1-treated mice (Fig. 4b, c, Supplementary Fig. 6d), indicating improved glucose-responsive serum insulin levels (Fig. 4d). Intraperitoneal glucose tolerance test (i.p.GTT) at 13 weeks of age further confirmed improved metabolic function in the Fex + JQ1-treated mice (Fig. 4e, f). Consistent with these findings, *ex vivo* GSIS assays demonstrated enhanced insulin secretion in isolated islets from Fex- and Fex + JQ1-treated mice (Fig. 4g). Notably, the combined treatment did not significantly affect body weight, food intake or insulin sensitivity measured by the i.p. insulin tolerance test (i.p.ITT), suggesting that glycemic improvement was due to enhanced β cell function rather than peripheral insulin sensitivity. (Supplementary Fig. 6d, e, f, Fig. 4h). Histological analysis revealed that Fex + JQ1 treatment significantly increased islet area and insulin-positive β cell mass compared to vehicle or single treatments (Fig. 4i, j, m), while reducing peri-islet fibrosis (Fig. 4i, k). The β-to-α cell ratio remained unchanged (Fig. 4l), indicating preservation of islet composition. These results demonstrate that Fex and JQ1 combined treatment improves islet mass and β cell function during the progression of T2D in *db/db* mice.

**Fig. 4.**
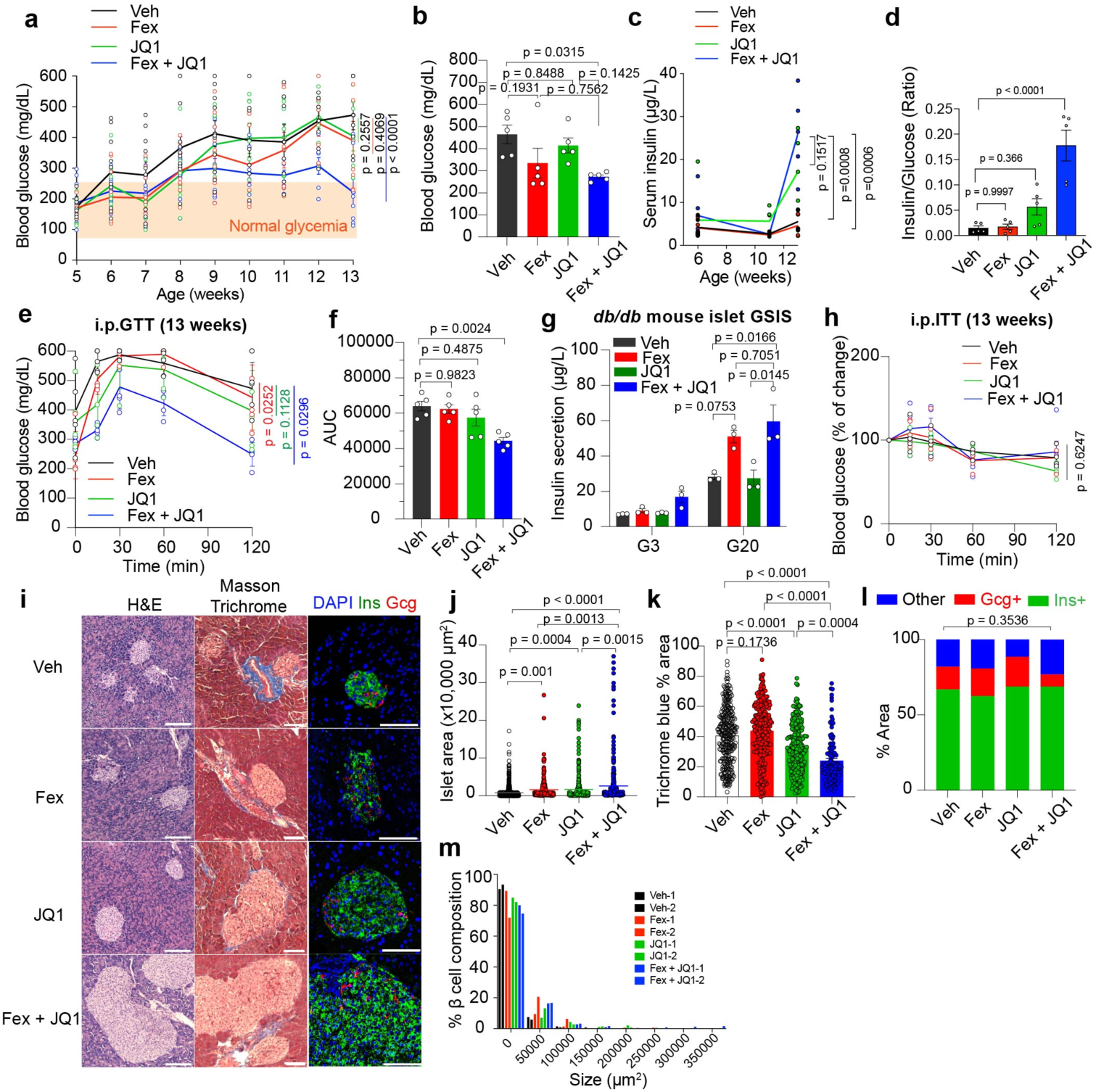
**FXR agonist and BET inhibitor improve islet function and glucose homeostasis in *db/db* mice**. **a**, Fasting blood glucose (mg/dL) of *db/db* mice during the 8-week course of i.p. treatment with vehicle (Veh), Fex, JQ1, Fex + JQ1 combination. n = 10 from 5 weeks to 9 weeks, and n = 7 from 10 weeks to 13 weeks (dissected n = 3/each condition at 9 weeks). Two independent cohort studies were merged. **b**, Fed *ad lib* blood glucose (mg/dL) of *db/db* mice on the 8-week of i.p. treatment with Veh, Fex, JQ1, Fex + JQ1 combination. N = 5/each. **c**, Serum insulin at three different time points of mice during the 8-week course of i.p. treatment with Veh, Fex, JQ1, Fex + JQ1 combination. n = 5/each. **d**, Serum insulin/glucose index at the on the 8-week of i.p. treatment with Veh, Fex, JQ1, Fex + JQ1 combination. n = 5/each. **e**, Intraperitoneal glucose tolerance test on the 8-week of i.p. treatment with Veh, Fex, JQ1, Fex + JQ1 combination. n = 5/each. **f**, Area under the curve (AUC) for the intraperitoneal glucose tolerance test at the 8-week of i.p. treatment with Veh, Fex, JQ1, Fex + JQ1 combination. n = 5/each. **g**, GSIS assay for the isolated islets from the *db/db* mice at the 8-week of i.p. treatment with Veh, Fex, JQ1, Fex + JQ1 combination. 10 islets/condition, n = 3/each. **h**, Insulin tolerance test blood glucose of *db/db* mice after 8-week i.p. treatment with Veh, Fex, JQ1, Fex + JQ1. n = 5/each. **i**, Representative images for H&E stain, Masson’s stain, and immunofluorescence stain for the pancreas of *db/db* mice after the 8-week of i.p. treatment with Veh, Fex, JQ1, Fex + JQ1 combination. Scale bar = 100 µm. **j**, Area of islets from H&E staining of *db/db* mouse pancreas after 8-weeks of i.p. treatment with vehicle, Fex, JQ1, Fex + JQ1 combination. Two histology slides per condition. **k**, Trichrome blue positive area from Masson’s trichome staining of *db/db* mouse pancreas after 8-week of i.p. treatment with Veh, Fex, JQ1, Fex + JQ1 combination. Two histology slides per condition. **l**, Percentage of insulin Alexa-488-positive and glucagon Alexa-647-positive areas of mouse islets from IHC staining of *db/db* mouse pancreas after 8-week of i.p. treatment with vehicle, Fex, JQ1, Fex + JQ1 combination. n = 10 islets. **m**, % of β cell composition. Statistical analyses were performed using two-way ANOVA for (a, c, e, h) and one-way ANOVA with Tukey’s multiple-comparison test for (b, d, f, g, j, k, l). Error bars represent ± SEM.

To understand the molecular basis of these effects, we performed transcriptomic analysis of isolated islets from Fex+JQ1 treated *db/db* mice. We identified 552 upregulated and 1,186 downregulated genes compared to Veh-treated controls (Fig. 5a, b). GO analyses revealed that pathways related to the functional β cells, such as the glucose response and insulin secretion, were upregulated (Fig. 5a). In contrast, the pathways related to the initiation of β cell dysfunction, such as the inflammatory response, response to hypoxia and regulation of the apoptotic process, were downregulated (Fig. 5b). Expression of key β cell identity markers (*Ins1*, *Ins2*, *Mafa*, and *G6pc2*) were elevated, especially in the combined treated group. Conversely, inflammatory genes (*Cxcl16*, *Tnfrsf1b*, *Tgfb2*, and *Cck*) were suppressed. We observed that similar pathways were respectively upregulated and downregulated in isolated islets from *db/db* mice treated with Fex or JQ1 alone (Supplementary Fig. 6g-j). We found that the key β cell functional and identity genes such as *Ins2*, *Ins1*, *Mafa* and *G6pc2* were upregulated in the isolated islets from Fex, JQ1 or Fex + JQ1 combined treated *db/db* mice, while the most profound upregulation was observed by Fex + JQ1 combined treatment (Fig. 5c). We found that the inflammatory pathway-related genes such as *Cxcl16*, *Tnfrsf1b*, *Tgfb2* and *Cck* were downregulated in the isolated islets from Fex, JQ1 or Fex + JQ1 combined treated *db/db* mice, while the most profound downregulation was observed upon Fex + JQ1 combined treatment (Fig. 5d). Motif analysis showed upregulated genes were associated with transcription factors (TFs) involved in insulin secretion and β cell differentiation, whereas downregulated genes were linked to inflammation and proliferation (Fig. 5e). In contrast, inflammation and fibrosis-related gene expression were not altered by the Fex and/or JQ1 treatment in the liver of *db/db* mice (Supplementary Fig. 6k). These findings suggest that the combined FXR and BET inhibition improves islet function by attenuating inflammation and preserving β cell identity *in vivo*. Given that JQ1 targets both BD1 and BD2 bromodomains of the BET protein, and recent studies suggest BD2-specific inhibitors regulate inflammation selectively^62^, we tested whether FXR activation combined with BD2-selective inhibition (BD2i/GSK620) recapitulates the benefits of JQ1 (Fig. 5f). Treatment with a BD2i in combination with Fex similarly improved fasting glucose levels and glucose tolerance (Fig. 5f–h, Supplementary Fig. 7a–e). Importantly, combined treatment with Fex + JQ1 or Fex + BD2i led to a beneficial reduction in serum cholesterol levels without significant alterations in other metabolic parameters, including aspartate aminotransferase (AST), alanine aminotransferase (ALT), amylase, creatinine, and triglycerides (Supplementary Fig. 8a–f). Given the well-established link between dyslipidemia and impaired glucose metabolism, these improvements in serum cholesterol may also contribute to the enhanced glucose tolerance observed in treated mice. Histological analysis of liver, kidney, intestine, and white adipose tissue (WAT) by H&E staining showed no detectable abnormalities (Supplementary Fig. 8g–j).

**Fig. 5.**
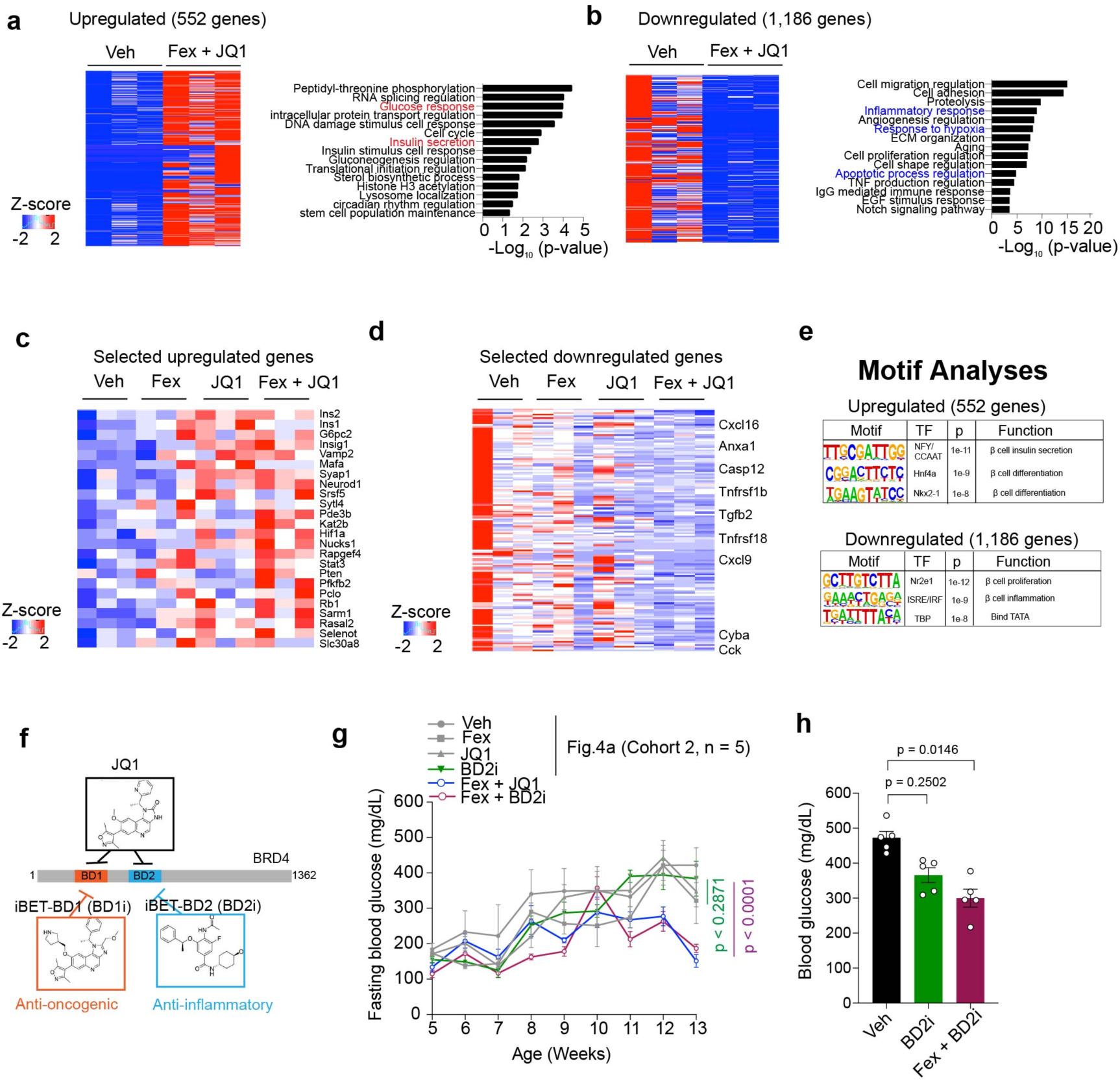
FXR agonist and BET inhibitor suppress inflammation-related gene expression. **a**, Upregulated genes and corresponding gene ontology pathway of islets from Veh-treated vs Fex+JQ1-treated *db/db* mice. n = 3/each. **b**, Downregulated genes and corresponding gene ontology pathway of islets from Veh-treated vs Fex + JQ1-treated *db/db* mice. n = 3/each**. c**, Heatmap of z-score of selected upregulated genes by Fex + JQ1 treatment related to glucose response and insulin secretion pathway. n = 3/each. **d**, Heatmap of z-score of selected downregulated genes by Fex + JQ1 treatment related to inflammation and apoptosis pathway. n = 3/each. **e**, Promoter motif analysis of the 552 upregulated genes or 1,186 downregulated genes by Fex + JQ1 treatment. **f**, Chemical structures and illustration of selective BD domain inhibitors, JQ1, BD1i and BD2i. **g**, Fasting blood glucose (mg/dL) of *db/db* mice during the 8-week i.p. treatment with BD2i single treatment and Fex + iBET-BD2 combination treatment. Gray bar shows Veh, Fex, JQ1, or Fex+JQ1 combination (Fig. 4a). Experiments were performed simultaneously with the experiments in Fig. 4a cohort 2. n = 5/each. **h**, Fed *ad lib* blood glucose (mg/dL) of *db/db* mice on the 12-week of i.p. treatment with BD2i vs Fex + BD2i. n = 5/each. Statistical analyses were performed using two-way ANOVA for (g) and one-way ANOVA with Tukey’s multiple-comparison test for (h). Error bars represent mean ± SEM.

To determine whether the observed glycemic improvements are dependent on β cell–intrinsic FXR signaling, we evaluated the effects of FXR activation and BET inhibition in an independent T2D model (HFD + MLD-STZ), which is characterized by β cell dysfunction. In wild-type (WT) mice, combined treatment with Fex and JQ1 significantly improved glycemic control (Fig. 6a–d). In contrast, these effects were completely abolished in β cell–specific Fxr knockout (βFxrKO) mice (Fig. 6e–h, Supplementary Fig. 9a, b), indicating that the metabolic benefits of Fex + JQ1 require β cell FXR. Consistently, both fed serum insulin levels and GSIS were increased in Fex + JQ1–treated WT mice (Fig. 6i, j), whereas these effects were absent in βFxrKO mice (Fig. 6k, l). Histological analysis further showed that increased islet mass and reduced fibrosis observed in WT mice were not present in βFxrKO mice (Fig. 6m–v). Together, these findings demonstrate that the protective effects of combined FXR activation and BET inhibition are dependent on β cell FXR signaling, establishing a direct mechanistic link between FXR activity and improved β cell function *in vivo*. To assess the relevance of this pathway in autoimmune T1D, we evaluated the effect of Fex and JQ1 in the spontaneous T1D model NOD mice. Fex alone delayed diabetes onset, with no additional benefit observed from the Fex + JQ1 combination at the tested dose. (Supplementary Fig. 9c–e). These results suggest that FXR activation is sufficient to confer protection in this context, whereas the contribution of BET inhibition may depend on disease context or require further optimization. Collectively, these findings identify β cell FXR as a critical mediator of metabolic protection and support a model in which BET signaling modulates, rather than independently drives, FXR-dependent β cell resilience *in vivo*.

**Fig. 6.**
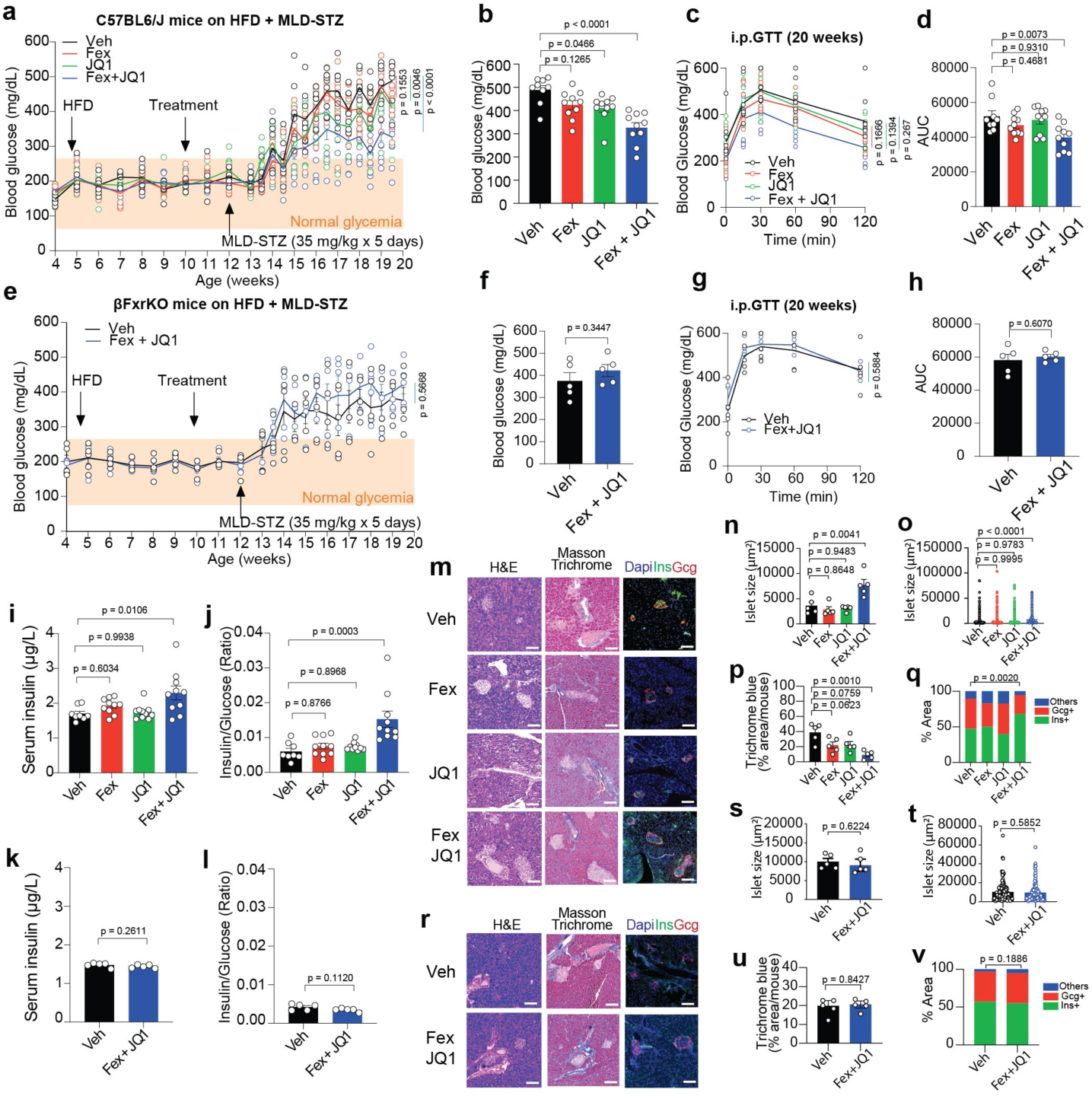
FXR agonism and BET inhibition improve glucose homeostasis in a HFD + MLD-STZ model, whereas these effects are abolished in β cell–specific Fxr knockout mice. **a**, WT (C57BL/6J) mice were fed a high-fat diet (HFD) and administered multiple low-dose STZ (MLD-STZ) for 5 consecutive days. Fed *ad lib* blood glucose levels (mg/dL) in WT mice treated with vehicle (Veh), Fex, JQ1, or Fex + JQ1 for 10 weeks. n = 9–10 per group. **b**, Fed *ad lib* blood glucose levels (mg/dL) in WT mice with or without HFD + MLD-STZ at 20 weeks of age. **c**, Intraperitoneal glucose tolerance test (i.p.GTT) in WT mice on HFD + MLD-STZ at 20 weeks of age after 10 weeks of treatment with Veh, Fex, JQ1, or Fex + JQ1. **d**, GTT area under the curve (AUC) in WT mice on HFD + MLD-STZ at 20 weeks of age after 10 weeks of treatment. **e**, *Ad lib* blood glucose levels (mg/dL) in β cell–specific Fxr knockout (βFxrKO) mice on HFD + MLD-STZ after 10 weeks of treatment with Veh or Fex + JQ1. n = 5 per group. **f**, Fed *ad lib* blood glucose levels (mg/dL) in βFxrKO mice on HFD + MLD-STZ at 20 weeks of age after 10 weeks of treatment. N = 5 per group. **g**, i.p.GTT in βFxrKO mice on HFD + MLD-STZ at 20 weeks of age after 10 weeks of treatment. n = 5 per group. **h**, GTT AUC in βFxrKO mice on HFD + MLD-STZ at 20 weeks of age after 10 weeks of treatment. n = 5 per group. **i**, Serum insulin levels in WT mice on HFD + MLD-STZ at 20 weeks of age after 10 weeks of treatment. **j**, Insulin-to-glucose ratio in WT mice under the same conditions. **k**, Serum insulin levels in βFxrKO mice on HFD + MLD-STZ at 20 weeks of age after 10 weeks of treatment. n = 5 per group. **l**, Insulin-to-glucose ratio in βFxrKO mice under the same conditions. n = 5 per group. **m**, Representative images of pancreatic islet histology from WT mice on HFD + MLD-STZ, including H&E staining, Masson’s trichrome staining, and immunofluorescence staining. Scale bar, 100 μm. **n, o**, Quantification of islet size from WT mice on HFD + MLD-STZ, shown as averages per mouse (n) and per group (o). Data were obtained from five independent mice per group. **p**, Quantification of trichrome-positive (blue) area per mouse from WT mice on HFD + MLD-STZ. **q**, Quantification of Insulin Alexa-488-positive and Glucagon Alexa-647-positive area composition in islets from WT mice on HFD + MLD-STZ. **r**, Representative images of pancreatic islet histology from βFxrKO mice on HFD + MLD-STZ, including H&E staining, Masson’s trichrome staining, and immunofluorescence staining. Scale bar, 100 μm. **s, t**, Quantification of islet size from βFxrKO mice on HFD + MLD-STZ, shown as averages per mouse (s) and per group (t). Data were obtained from five independent mice per group. **u**, Quantification of trichrome-positive (blue) area per mouse from βFxrKO mice on HFD + MLD-STZ. **v**, Quantification of Insulin Alexa-488-positive and Glucagon Alexa-647-positive area composition in islets from βFxrKO mice on HFD + MLD-STZ. Statistical analyses were performed using two-way ANOVA for (a, c, e, g), one-way ANOVA with Tukey’s multiple-comparison test for (b, d, i, j, n, o, p, q), and unpaired two-tailed Student’s t-test for (f, h, k, l, s, t, u, v). Error bars represent mean ± SEM.

### FXR agonist and BET inhibitor protect human T2D and T1D models *in vitro*

One of the major challenges of drug development is the limited translational relevance of preclinical models to human disease. Although *in vivo* mouse models are valuable for screening drug candidates, many compounds fail to translate effectively to the human clinical setting^63^. To assess the clinical relevance of FXR–BET targeting, we employed human pluripotent stem cell–derived islet-like organoids (HILOs) as *in vitro* models of human T2D and T1D^46,64^ (Fig. 7a, Supplementary Fig. 10a). Our previous studies have shown that thioredoxin-interacting protein (TXNIP) plays a key role in the induction of β cell dysfunction^65^. It is known that human but not rodent islet amyloid polypeptide (IAPP) accumulation and elevation of TXNIP lead to ER stress-induced β cell apoptosis in T2D^65–68^. Similarly, human IAPP and TXNIP lead to NLRP3 inflammasome activation to trigger IL-1β and CASP1 activation, providing a mechanism of β cell dysfunction in T2D islets^69,70^.

**Fig. 7.**
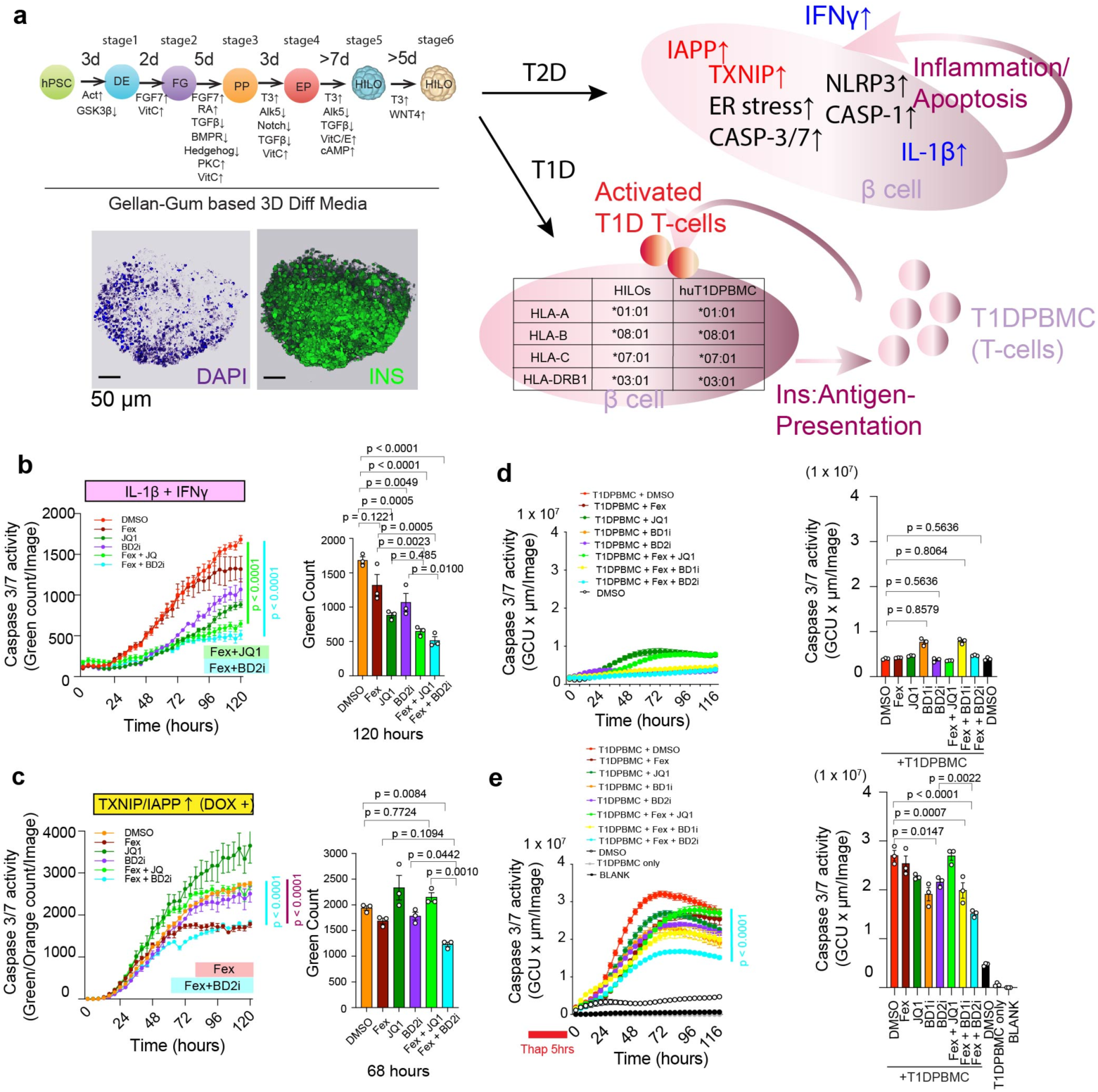
FXR agonist and BET inhibitor protect human iPSC-derived human islet-like organoid models of human T1D and T2D *in vitro.* **a**, Schematic of the differentiation strategy into HILOs. IHC of insulin Alexa-488 and DAPI of day 30 HILOs are shown. Illustration of the protocol for iPSC-derived human islet-like organoid (HILO) *in vitro* T1D and T2D models. T2D includes HILO with DOX-induced overexpression of human IAPP and TXNIP. The T1D HILO model is the co-incubation of HILO with peripheral blood mononuclear cells carrying the matched with HLA-A*01:01 HLA-B*08:01, HLA-C*07:01 HLA-DRB1*03:01 as the HILO. Scheme of partially HLA-matching was shown. **b**, Apoptosis rate in dispersed HILOs was measured by real-time imaging using Incucyte SX5. Dispersed HILOs were treated with DMSO, Fex (10 µM), JQ1 (500 nM), BD2i (1 µM), Fex (10 µM) + JQ1 (500 nM), or Fex (10 µM) + BD2i (1 µM), in the presence of IL-1β (10 ng/ml) and IFNγ (10 ng/ml). n = 3/each. Apoptosis was visualized by Caspase 3/7 green fluorescence. Bar graph shows green (apoptosis) cell count at 120 hours. **c**, Caspase 3/7 green (Green = Caspase 3/7 activity/Orange = human TXNIP/IAPPOE cells count) of T2D HILO model (DOX-inducible human TXNIP and human IAPP inducing) during 120 hours treatment of DMSO, Fex (10 µM), JQ1 (500 nM), BD2i (1 µM), Fex (10 µM) + JQ1 (500 nM), or Fex (10 µM) + BD2i (1 µM). n = 3/each. **d**, Apoptosis rate in dispersed HILOs was measured by real-time imaging using Incucyte SX5. Dispersed HILOs were co-cultured with human T1DPBMC and treated with Fex (10 µM), JQ1 (500 nM), BD1i (1 µM), or BD2i (1 µM) with or without IFNγ (10 ng/ml) for 120 hours. n = 3/each. **e**, Apoptosis rate in dispersed HILOs was measured by real-time imaging using Incucyte SX5. Dispersed HILOs were co-cultured with human T1DPBMC and treated with Fex (10 µM), JQ1 (500 nM), BD1i (1 µM), or BD2i (1 µM) with or without IFNγ (10 ng/ml) for 120 hours. Thap (Thapsigargin, ER stress inducer, 5 µM) were pretreated to enhance the expression of misfolded insulin prior to imaging. n = 3/each. Statistical analyses were performed using two-way ANOVA for the left panels of (b–e) and one-way ANOVA with Tukey’s multiple-comparison test for the right panels of (b–e). Error bars represent mean ± SEM.

We created an *in vitro* human T2D model system by establishing doxycycline (DOX)-inducible human IAPP and TXNIP in human induced pluripotent stem cells (hiPSCs) and differentiating them into HILOs (hereafter referred to as *in vitro* HuT2D model system/ivHuT2D). In this model, we observed that the simultaneous induction of TXNIP and/or IAPP led to an increase in PERK, cleaved caspase 3, and cleaved IL-1β expression in β-like cells (Supplementary Fig. 10b). Using real-time imaging, we determined apoptosis by co-induction of IAPP and TXNIP in β-like cells from dispersed HILOs. On day 30, HILOs were dispersed in two-dimensional (2D) culture and treated with DOX and/or IL-1β + IFNγ in the presence of DMSO, Fex, JQ1, Fex + JQ1, BD2i or Fex + BD2i. We found that both IL-1β + IFNγ treatment and co-induction of IAPP and TXNIP induced caspase 3 activation in β-like cells from dispersed HILOs (Fig. 7b, c). Live cell imaging analyses revealed that the apoptosis induced by both IL-1β + IFNγ treatment and DOX-dependent co-induction of IAPP and TXNIP was significantly suppressed by Fex + BD2i and/or Fex + JQ1 treatment in β-like cells (Figs. 7b, c, Supplementary Fig. 10c, d). These results suggest that FXR and BET signaling can improve human IAPP and TXNIP-induced β cell toxicity in ivHuT2D.

Next, to test whether similar efficacy could be observed in an antigen-mediated autoimmune T1D model, we established the human T1D response *in vitro* by co-culturing partially MHC-matched dispersed HILOs (HLA-A*01:01 HLA-B*08:01, HLA-C*07:01 HLA-DRB1*03:01) and peripheral blood mononuclear cells (PBMCs) derived from human T1D patients (HLA-A*01:01 HLA-B*08:01 HLA-C07:01 HLA-DRB1*03:01, HemaCare) (hereafter referred as *in vitro* HuT1D model system/ivHuT1D). We determined the apoptosis induction rate in HILOs by insulin antigen-responsive T-cells in PBMCs. As an enhanced immune response, PBMCs were stimulated with anti-CD3/CD28 and HILOs were pretreated with thapsigargin (Thap, enhanced by prior ER-stress induction and IFNγ treatment^71^) for 5 hours to increase the expression of misfolded insulin. Consistent with previous reports, we observed that pretreatment of Thap significantly enhanced T1DPBMC mediated apoptosis in HILOs (Fig. 7d, e). We co-treated these cells with Fex, JQ1 and iBET-BD1 (BD1i) or BD2i and found that T1DPBMC promoted apoptosis in MHC-partially matched HILOs (Fig. 7d, e). Importantly, Fex combined with BD2i exhibited the strongest protective effect, suggesting that selective inhibition of the BD2 domain enhances the anti-inflammatory effects of FXR activation in human β cells (Fig. 7e). Collectively, these findings demonstrate that FXR activation and BET inhibition cooperatively protect human β cells in both T2D and T1D-relevant contexts, highlighting the translational potential of targeting the FXR–BET axis.

## Discussion

Here, we identify a BA–epigenetic regulatory axis in which FXR and BRD4 cooperatively control β cell inflammatory responses and functional identity during diabetes progression. Current diabetic treatments primarily focus on enhancing insulin secretion or function. However, few therapies exist to prevent the loss and preserve the functional mass of β cells in patients with T1D and T2D. Notably, the onset of hyperglycemia in T1D and T2D typically requires the destruction of more than ∼50% of β cells^4^. Thus, therapeutic strategies that actively protect and preserve β cell function represent an urgent unmet clinical need. This work builds upon growing evidence that BET family proteins regulate NR signaling. BRD9 has been shown to interact with the vitamin D receptor (VDR)^54^, repressing its anti-inflammatory actions in β cells, while also associating with the glucocorticoid receptor (GR) to limit anti-inflammatory responses^57^. In parallel, BRD4 facilitates transcription by recognizing acetylated histones and non-histone proteins, thereby influencing both inflammation and cell fate. FXR, conversely, acts as a transcriptional repressor under inflammatory stress by recruiting the Silencing Mediator for Retinoid and Thyroid hormone receptors (SMRT) corepressors^72^, and is known to regulate insulin secretion through genomic and non-genomic pathways. We now extend this paradigm by demonstrating that FXR acetylation at K158/K218 is essential for its interaction with BRD4, and that this interaction dynamically regulates key β cell transcriptome and function in disease conditions. BA pathways play a key role in physiological regulation of β cell function, identity, and integrity. Our results demonstrate that dysregulated BA signaling actively contributes to β cell vulnerability by accelerating cytokine-induced dysfunction in both mouse and human β cells. Our findings demonstrate clear therapeutic potential, as shown by simultaneous targeting of BA sensor FXR signaling and the epigenetic reader BET signaling that synergistically improved β cell function in our *in vivo* and *in vitro* models. These findings establish FXR–BET signaling as a central regulatory axis governing β cell inflammatory resilience and identity.

BRD4 is a known regulator of inflammatory responses and cell fate-determination by binding to the acetylated histones and non-histone proteins, a marker of active transcription to facilitate the recruitment of positive transcription elongation factors^38^. While FXR–BRD4 synergy has been explored in the context of hepatic injury, recent reports have also highlighted that co-treatment with the FXR agonist obeticholic acid (OCA) and BRD4 inhibitor JQ1 can paradoxically exacerbate liver inflammation and fibrosis under certain cholestatic conditions, suggesting context-specific adverse effects^42^. In contrast, our findings uncover a novel and tissue-selective mechanism within pancreatic islets, where co-targeting FXR and BRD4 exhibits protective effects. When mice with α-naphthylisothiocyanate (ANIT)-induced liver injury were treated with the endogenous FXR agonist OCA and JQ1 individually via oral gavage over seven days, significant improvements in hepatotoxicity, liver inflammation, and fibrosis were noted^42^. Diverging from this gut-targeted approach, our study implemented an intraperitoneal treatment regimen of Fex and JQ1, three times weekly over eight weeks, which was more effective in preserving β cell function and improving glycemic control in *db/db* mice (Fig. 4a-g). Unlike OCA, Fex is a synthetic FXR agonist that reaches the pancreas via intraperitoneal injection and is not subject to microbial conversion into secondary BAs, potentially contributing to its enhanced efficacy and reduced systemic toxicity. Interestingly, while Fex and JQ1’s synergy was evident in islets, it was not observed in the liver (Supplementary Fig. 6k), underscoring the tissue-specific nature of FXR and BET signaling. These results highlight that the therapeutic efficacy of FXR–BET targeting is highly dependent on ligand pharmacology, delivery route, and tissue-specific regulatory context. Aligned with this, we showed the combined influence of an FXR agonist and the BRD4 inhibitor JQ1 in dynamically modulating inflammatory genes and β cell identity genes under cytokine-induced stress in EndoC-βH1 cells (Fig. 3f, Supplementary Fig. 4e) and during the evolution of the diabetes in *db/db* islets (Fig. 5c, d). These observations imply that FXR and BRD4 mutually fine-tune gene expression to regulate physiological responses in β cells. Additionally, studies have also highlighted the role of BRD4 in pancreatic development by regulating the expression of fate-determinant factors^73^ and its inhibitory role in insulin transcription and secretion in β cells^44^. In contrast, FXR positively regulates insulin transcriptome and secretion in β cells^25,26^. Consistent with this reciprocal regulation by FXR and BRD4 on β cell identity and function, we noticed a synergistic enhancement in the expression of critical β cell identity genes, *Ins2*, *G6pc2*, and *Neurod1* in the presence of Fex and JQ1 (Fig. 5c). Specifically, our data indicate that the acetylation of K158/K218 may be essential for BRD4 recruitment (Fig. 2h). It is established that FXR undergoes acetylation by P300 and deacetylation by Sirtuin 1 (SIRT1), influencing its stability and its ability to form heterodimers with Retinoid X Receptor α (RXRα)^58^. Further investigations are warranted to clarify how these post-translational modifications influence BRD4 binding to FXR and how this interaction impacts β cell identity and function.

A well-recognized limitation of preclinical drug development is the frequent discrepancy between therapeutic efficacy observed in mouse models and poor translation to human patients. For example, the reported beneficial effects of TUDCA are largely confined to conditions of severe endoplasmic reticulum (ER) stress, such as high-dose TUDCA (>100 μM) in combination with thapsigargin, a potent ER stress inducer, in porcine islets *in vitro*^74^ or in STZ-induced diabetic mice *in vivo*^75^. In contrast, our data indicate that TUDCA functions as a competing agonist or functional antagonist of CA, thereby suppressing FXR activation in human β cells. This divergence likely reflects intrinsic species-specific differences and context-dependent effects in BA receptor signaling. Recent evolution of stem cell-derived organoid technology provides a window into how human organs might respond *in vitro*^76^. Our application of such models to this study showed that when *TXNIP* and *IAPP* were forcefully introduced into HILOs, it emulated the induction of apoptosis by increasing ER-stress and caspase activation observed in human β cells affected by T2D (Supplementary Fig. 10b, c). Similarly, using a recently introduced *in vitro* platform in modeling characteristics of human T1D by utilizing stress-induced cells derived from patients, paired with their native immune cells, successfully recreates the autoimmune reaction seen in T1D^71^. Our studies indicate that co-culturing partially MHC-matched, stress-induced HILOs with PBMC from T1D patients leads to an autoimmune-induced apoptosis (Fig. 7e). In this setting, combined treatment with Fex and JQ1 also augmented the survival rate of dispersed HILOs. Importantly, these platforms provide a tractable model for dissecting human-specific immunometabolic interactions and therapeutic responses.

Clinically, our findings carry significant translational implications. By targeting both metabolic (FXR) and epigenetic (BRD4) regulators, this combined strategy offers a pathway to restore endogenous insulin production in both T1D and T2D, independent of immune modulation or β cell replacement. Moreover, BD2-selective BET inhibitors, which are in early-phase trials for inflammatory diseases, may offer a favorable safety profile in combination therapies. This work may guide future precision medicine approaches where BA profiling and epigenetic signatures could stratify patients most likely to benefit from FXR–BET modulation.

In summary, our study uncovers a previously unrecognized BA–epigenetic axis that integrates metabolic and chromatin signaling to control β cell fate under stress. By demonstrating that FXR activation and BET inhibition cooperatively preserve β cell function across mouse and human systems, our work establishes a mechanistic and translational foundation for β cell–protective therapies in diabetes.

### Limitations of the study

While our study demonstrates the therapeutic potential of FXR activation combined with BET inhibition in both mouse and human models of diabetes, several limitations should be acknowledged. First, the dosing regimen (50 mg/kg Fex and 25 mg/kg JQ1 administered intraperitoneally three times per week) was optimized for proof-of-concept efficacy. Further dose–response studies, alternative routes of administration, and long-term treatment analyses will be required to refine therapeutic windows and assess safety profiles. Second, although certain experiments (e.g., histological analyses) were performed with relatively limited sample sizes due to the breadth of multi-level analyses (*in vivo*, *ex vivo* islet assays, and transcriptomics), the key findings were consistently reproduced across multiple independent experimental systems, including mouse models, primary islets, and human β cell platforms. In addition, we acknowledge that differences in cytokine combinations, incubation conditions, and functional readouts across cell systems may introduce variability in the magnitude of observed effects. These differences primarily reflect intrinsic physiological properties of each model (e.g., limited glucose responsiveness in EndoC-βH1 cells) rather than experimental inconsistency. Notably, when key experimental parameters were harmonized, the synergistic effects of FXR activation and BET inhibition were consistently observed across systems, supporting the robustness of the central conclusions. Third, while our human iPSC-derived islet organoid (HILO) models recapitulate key features of β cell dysfunction in T1D and T2D, they do not fully capture the chronic and multifactorial nature of disease progression *in vivo*. In particular, T1D pathogenesis involves complex and evolving immune interactions, including antigen spreading, epitope diversification, and regulatory immune networks, which are difficult to model in short-term co-culture systems. Our current model primarily reflects β cell–T cell interactions and may underrepresent contributions from additional immune components such as B cells, macrophages, and regulatory T cells, as well as systemic inflammatory cues. Future studies incorporating more complex immune environments and longer-term culture systems will be necessary to improve translational relevance. Finally, FXR and BRD4 are known to have stage-specific roles during development and differentiation. As our current analyses are focused on mature β cells under stress conditions, it remains unclear whether FXR–BET interactions similarly regulate earlier stages of islet development or maturation. Future studies using temporally controlled genetic models or developmental organoid systems will be required to address these questions. Overall, these limitations highlight areas for further investigation while supporting the central conclusion that FXR-dependent signaling plays a key role in β cell protection under metabolic and inflammatory stress.

## METHOD DETAILS

### Resources

All key resources used this study are listed in Supplementary Table.3.

### Mouse line maintenance

Animals were maintained in a pathogen-free animal facility on a 12-hour light/dark cycle at an ambient temperature of 23°C. Water and food were provided *ad lib*. C57BL/6J (JAX000664) and BKS.Cg-Dock7m +/+ Leprdb/J (*db/db*, JAX000642) were purchased from The Jackson Laboratory. Fxr flox/flox mice^59^ were obtained from Dr. Thomas Q. de Aguiar Vallim, and Brd4 flox/flox^42^ mice from Dr. Lin-Feng Chen. Mice were injected intraperitoneally three times a week as a single treatment or in combination with 50 mg/kg Fexaramine (Fex, AdooQ Bioscience, A21771), 25 mg/kg JQ1 (AdooQ Bioscience, A12729), 10 mg/kg GSK620 (iBET-BD2, AdooQ Bioscience, A22521), or vehicle (0.9% saline solution, Growcells, MSDW1000) containing 10% 2-hydroxypropyl-β-cyclodextrin (Sigma H-107). Blood glucose levels were measured weekly in both fed *ad lib* and 6-hour-fasted animals. Blood was collected via tail-prick, and glucose concentrations were determined using NovaMax glucose test strips (Nova Medical, REF43523) and a NovaMax Plus glucose meter (Nova Medical, REF8548043435). All animal procedures were performed in accordance with IACUC approved protocols (#32082-03) and animal resources at the Lundquist Institute for Biomedical Innovations at UCLA-Harbor Medical.

### Intraperitoneal glucose tolerance test and insulin tolerance test

Mice were fasted for 12-16 hours prior to intraperitoneal injection of glucose. 1 g of glucose per kilogram (kg) of body weight dissolved in 0.9% saline was injected intraperitoneally, and blood glucose was measured at time 0, 15, 30, 60, and 120 minutes after injection. For the insulin tolerance test, mice were fasted for 4 hours prior to the insulin injection. Humulin R U-100 insulin (Eli Lilly, HI-210) was diluted in 0.9% saline solution (Growcells, MSDW1000) and then injected intraperitoneally at 0.75 unit per kg of body weight, and blood glucose was measured via tail prick using NovaMax glucose strip (Nova Medical, 43523) and NovaMax Plus glucose meter (Nova Medical, 8548043435) at 0, 15, 30, 60, and 120 minutes after injection.

### Serum bile acid profiling

The C57BL/6J (male, 8 weeks), BKS (male, 8 weeks), *db/db* (male, 8 weeks), NOD (female, 19 weeks, diabetic and non-diabetic) and NOD (female, 8 weeks and 23 weeks) mice were sacrificed according to the IACUC approved protocol. As BA profile can be affected by multiple factors, mice were euthanized under similar conditions - namely, mice were kept in the same facility, euthanized in the same facility with the same equipment, at the similar time of day without fasting at approximately 13:00 PST, limited to 10 mice per day, with collection performed on successive days for groups that were compared with each other. Blood was collected in BD Microtainer® blood collection tubes (BD microtainer # 365985) and stored overnight at 4°C prior to centrifugation at 3000 g for 15 minutes. The serum fraction was collected and stored at -80°C. UPLC separation of bile acids was performed using a Waters (Milford, MA) Acquity UPLC with a proprietary reverse phase UPLC column and guard column provided by Biocrates. The analytes were separated using a gradient of 10 mm ammonium acetate, 0.015% formic acid in water to 10 mm ammonium acetate, 0.015% formic acid in acetonitrile (65%) and methanol (30%). Total UPLC analysis time was approximately 6 minutes per sample. Using electrospray ionization in negative ion mode, samples were introduced directly into a Xevo TQ-S triple quadrupole mass spectrometer (Waters) operating in the multiple reaction monitoring (MRM) mode. MRM transitions (compound specific precursor to product ion transitions) for each analyte and internal standard were collected over the appropriate retention time. The UPLC-MS/MS data were imported into the Waters TargetLynx™ application for peak integration, calibration, and concentration calculations. The UPLC-MS/MS data from TargetLynx™ were analyzed using Biocrates Me*tIDQ*™ software. Bile acid profiling was performed at the Duke University Proteomics and Metabolomics Core Facility.

### Serum insulin and Metabolites

Serum insulin was measured using a mouse insulin ELISA kit (Mercodia, 10-1247-01) according to the manufacturer’s protocol. Serum metabolites were analyzed by IDEXX bioanalytic service.

### Histology, immunohistochemistry, microscopy and imaging analyses

H&E and Masson’s trichrome staining were performed on paraffin-embedded sections of mouse pancreas and visualized using the EVOS M7000 imaging system or Keyence BZ-X800. For immunofluorescence (IF) staining, paraffin-embedded sections of mouse pancreas were deparaffinized with xylene solution and rehydrated through a graded series of decreasing concentrations of ethanol. Antigen retrieval was performed at 85 – 115°C for 10 min in high pressure (0.2–11.6 psi) using antigen unmasking solution (Vector Laboratories, H-3300-250). Incubation with antibodies to insulin (1/100, Abcam, ab7842), glucagon (1/100, Abcam ab10988), somatostatin (1/100, Abcam, ab103790), with overnight incubation at 4°C. Secondary antibodies were coupled to Alexa Fluor 488 goat anti-guinea pig IgG or Alexa Fluor 647 donkey anti-mouse IgG (Life Technologies, A11073) and visualized by fluorescence confocal microscopy (Leica SP8 or Nikon Ti2 or Keyence BZ-X800). DAPI (VECTOR, H-1200) or Hoechst 33342 (Thermo Scientific, 2249, 1 µg/ml final concentration) was used for nuclear staining. Islet area quantification and image analyses were performed using ImageJ. Masson’s trichrome staining was quantified using the ImageJ Colour Deconvolution 2 plugin.

### Cell Culture and Treatment

EndoC-βH1 cells (UniverCell) were maintained in Cultrex RGF Basement Membrane Extract (Cultrex gel)-coated plates with EndoC-βH1 culture medium containing the following ingredients: MCDB131 medium (Sigma, M8537-10X1L), 0.1745% NaHCO_3_ (Spectrum, S1147), 2% free fatty acid-free BSA fraction V (FF-BSA, GoldBio, A-421-1), 1% Glutamax-I (Gibco, 35050-061), 1% penicillin/streptomycin (Gibco, 15140-122), 0.25 mM ascorbic acid (Sigma, A4544-25G), 10 mM zinc sulfate (Sigma, Z0251-100G), 55 µM 2-mercaptoethanol (Gibco, 21985-023), 1% MEM-NEAA (Gibco, 11140-050), 10 mM HEPES (Gibco, 15630-080), and 11 mM glucose (Fisher, BP350-1).

INS-1 cells were cultured at 37°C with 5% CO_2_ in air in RPMI1640 (Sigma, 11875-093) supplemented with 10% (v/v) fetal bovine serum (Genesee Scientific, 25-525H), 1% (v/v) penicillin/streptomycin antibiotics, 10 mM HEPES, 2 mM Glutamax-I, 1 mM sodium pyruvate and 55 µM 2-mercaptoethanol. EndoC-βH1 cells were passaged 1-2 times per week. INS-1 cells were passaged 2-3 times per week.

HEK293LTV cells (Cell Biolabs, LTV-100) were cultured in DMEM (Sigma, D6546-500ML) containing 10% FBS (Genesee Scientific, 25-525H), 1% Glutamax-I, and 1% penicillin/streptomycin. HEK293LTV cells were passaged every 2 days.

Human pluripotent stem cells (hPSCs), such as human induced pluripotent stem cells (ChiPSC12, TAKARA) were maintained in Cultrex RGF Basement Membrane Extract (Bio-techne, 3433-010-01)-coated plates with mTeSR Plus (STEMCell Technologies, 100-027). mTeSR Plus was replaced to fresh every other day. hPSCs were passaged every 4-7 days with ReLeSR (STEMCell Technologies, 05872). Single cell suspensions were prepared with Accutase (STEMCell Technologies, 07920), washed in PBS, and collected by centrifugation (1000-1300 rpm for 5 min).

hPSCs, EndoC-βH1 cells, and HEK293LTV cells were cryopreserved using Bambanker (FujiFilm Wako, 302-14681) in -80°C deep freezer or liquid nitrogen. Supplements of the following small molecule agonists and antagonists were prepared in dimethyl sulfoxide DMSO: Fex (10 μM), JQ1 (500 nM), GSK778 (1 μM, iBET-BD1, AOBIOUS, AOB11459), GSK620 (1 μM). Cells were pretreated with the small molecules for 24 hours prior to the addition of the above cytokines: 10 ng/ml IL-1β and 10 ng/ml IFNβ or 10 ng/ml IFNγ.

### Generation of human islet-like organoid (HILO) cultures

Human islet-like organoids (HILOs) were generated according to a previous study^46^ with minor modifications. Briefly, hPSCs were cultured in Cultrex gel-coated plates until it reaches to approximately ∼90% confluence. hPSC suspensions were prepared using ReLeSR or Accutase, and cultured in ultra-low attached 6-well plates (2 × 10^6^ cells/mL) with orbital shaking (∼60 rpm for 2 days), or EZSPHERE (IWAKI, 4810-900SP) or Agreewell 800 (StemCell Technologies, 34860) at 5,000-10,000 desired cell number per microwell in the presence of the ROCK inhibitor (10 μM Y-27632, StemCell Technologies, 72308). Cells were then resuspended with 3D Kelco Gel Stem TeSR™ Base Medium in the presence of the ROCK inhibitor (10 μM) for 0-2 days until spheroids reached ∼200 mm in diameter. The medium was then replaced with 0.015% Kelco gel containing 0.3% methylcellulose and supplemented with 100 ng/ml human Activin A and 3 μM CHIR99021 in differentiation medium (S1) for 1 day and then 100 ng/ml human Activin in differentiation medium (S1) for another 2 days (stage 1, definitive endoderm). Subsequently, the medium was changed to differentiation medium (S2) with 50 ng/ml FGF7 for 2 days (stage 2, foregut/FG), then to differentiation medium (S3) with 50 ng/ml FGF7, 0.25 μM SANT-1, 1 μM retinoic acid, 100 nM LDN193189, 10 μM Alk5 inhibitor II, and 200 nM of the α-amyloid precursor protein modulator TPB for 3 days (stage 3, pancreatic progenitor/PP), then 50 ng/ml FGF7, 0.25 μM SANT-1, 1 μM retinoic acid, 100 μM LDN193189, 10 μM Alk5 inhibitor II, and 100 nM of the α-amyloid precursor protein modulator TPB for 2 days. Subsequently, medium was replaced with differentiation medium (S4) with 0.25 μM SANT-1, 50 nM retinoic acid, 100 nM LDN193189, 10 μM Alk5 inhibitor II, 1 μM T3 for 3 days (stage 4, endocrine progenitor/EP). Subsequently, the medium was replaced with differentiation medium (S5) containing 100 nM LDN193189, 100 nM γ-secretase inhibitor XX, 10 μM Alk5 inhibitor II, and 1 μM T3 for 7 days (stage 5, immature β-like cells/imβ). Subsequently, the medium was replaced with differentiation medium (S5) containing 10 μM Trolox, 2 μM R428, 1 mM N-acetyl cysteine, 10 μM Alk5 inhibitor II, and 1 μM T3 for an additional 5 days (day 25) (stage 6, immature β-like cells/imβ). From day 26, the medium was replaced with differentiation media (S5) with 1 μM T3, 10 μM Alk5 inhibitor II and WNT4 supernatant (Final volume 10%, ∼100 ng WNT4) (stage 7, mature β-like cells/β). WNT4 supernatant was generated by DOX-inducible WNT4 expressing HEK293LTV. After 24 hours of DOX treatment, medium was replaced to S5 Media and subsequentially stimulated with DOX for 3–4 days. Supernatant was then filtered using 0.22 mm filter and the batch containing >100 ng/ml of WNT4 was frozen as stock until further use. After 30 days, the HILOs were maintained in differentiation media (S6) containing 1 μM T3. HILOs were dissociated into single cells by TrypLE with 10 ng/ml DNase I for 12 min at 37 °C shaking water bath and then cryopreserved using Bambanker at 10-20 million cells/ml until it is used for experiments.

For modeling human T2D *in vitro* (ivHuT2D), DOX-inducible human IAPP (dhIAPPOE) and human TXNIP (dhTXNIPOE) overexpression ChiPSC12 was established. ChiPSC12 derived HILOs (dhIAPPOE/dhTXNIPOE) were dispersed in Cultrex gel-coated 96 well plates at 100,000 cells/well, treated with DMSO, 10 μμ Fex, 500 nM JQ1, 1 μμ iBET-BD2 or combination of Fex and JQ or iBET-BD2 (GSK620) for 24 hours before being treated with 10 ng/ml IL-1β and 10 ng/ml IFNγ. For modeling human T1D *in vitro* (ivHuT1D) ChiPSC12 derived HILOs (HLA-A*01:01 HLA-B*08:01, HLA-C*07:01 HLA-DRB1*03:01) were dispersed in Cultrex gel-coated 96 well plates at 100,000 cells/well and treated with 5 μM thapsigargin (Sigma, T9033-1MG) for 5 hours and then the peripheral blood mononuclear cells (PBMCs) derived from human T1D patients (HLA-A*01:01 HLA-B*08:01 HLA-C07:01 HLA-DRB1*03:01, HemaCare) were added at 10,000 cells/well with 10 ng/ml IL-1β and 10 ng/ml IFNγ. Apoptosis induction in ivHuT2D and ivHuT1D was measured by Incucyte SX5 (Sartorius) with Incucyte Caspase-3/7 Green Dye (Sartorius, 4440).

### Mouse islet isolation

Mouse pancreatic islets were isolated as previously described^46,65^. Briefly, 0.5 mg/mL collagenase P (Roche, 11213873001) was dissolved in HBSS buffer (Gibco, 14170112) and injected into the common bile duct. The perfused pancreas was collected and incubated in a 37°C water bath with shaking for 21 min. Digested pancreatic exocrine tissue was subjected to density gradient centrifugation using Histopaque-1077 (Sigma H8889) at 900 × g for 15 min to separate exocrine tissue from intact islets. Islets were handpicked under a stereomicroscope and cultured in RPMI 1640 medium (Gibco, 11875-093) supplemented with 10% fetal bovine serum (FetalPURE; GenClone, 25-525H), 1% penicillin–streptomycin (Gibco, 15070-063), and 1% GlutaMAX (Gibco, 3505-061) for 24 hours prior to treatment. Isolated islets were treated with 10 μM Fex (AdooQ Bioscience, A21771), 500 nM JQ1 (AdooQ Bioscience, A12729), or 1 μM BD2-selective inhibitor (iBET-BD2/GSK620; AdooQ Bioscience, A22521), alone or in combination, for 24 hours, followed by cytokine stimulation with 10 ng/mL IFNγ (PeproTech, 300-02-100UG) and 10 ng/mL IL-1β (PeproTech, 200-01B-10UG) for an additional 48 hours. Human islets were precultured in CMRL 1066 or RPMI 1640 medium supplemented with 10% FBS, 1% ITS, 1% HEPES, 1% GlutaMAX, and 1% penicillin–streptomycin. Human islets were obtained from the Integrated Islet Distribution Program (IIDP; SAMN27619977, SAMN34411471, SAMN35848421) and subjected to the same treatment conditions as mouse islets unless otherwise indicated.

### Virus production

Lentiviruses were produced using second- or third-generation lentiviral systems in the HEK293LTV cell line. Briefly, viral core plasmids and target lentivirus plasmids were transfected into 70% confluent HEK293LTV cells using Lipofectamine 3000 (Invitrogen, L3000015) and Opti-MEM (Gibco, 31985070). The medium was changed to fresh HEK293LTV culture medium one day after transfection as day 0. Lentivirus-containing supernatants were collected on day 1 and day 3 and centrifuged at 1300 rpm for 5 minutes to remove any cell debris. The supernatants were further filtered through 0.45 µm filter, and the lentivirus was used as fresh virus or concentrated 10 times with DMEM/F12 using a Lenti X Concentrator (TAKARA, 631232) for long-term storage at -80°C in a deep freezer. 10 μg/ml Polybrene (Santa Cruz, sc-134220) was used as a scaffold for virus infection.

### GSIS assay

Isolated pancreatic islets were pre-cultured at 37°C for 30 min in Krebs Ringer Bicarbonate Buffer with HEPES (KRBH) containing 129.4 mM NaCl (Sigma, S7653-250G), 3.7 mM KCl (Fisher, BP366-1), 2.7 mM CaCl_2_ (Alfa Aesar, L13191), 1.3 mM KH_2_PO_4_ (Sigma, P5655-100G), 1.3 mM MgSO_4_ (Sigma, 63138-250G), 24.8 mM NaHCO_3_ (Sigma, 63138-250G), 10 mM HEPES (Bioland Scientific, CH01-500G) and 0.2% (v/v) FAF-BSA fraction V (GoldBio, A-421-1) with 3 mM glucose (Fisher, BP366-1). The islets were then incubated for 30 minutes in KRBH buffer (500 μl per 10 islets) containing 3 mM glucose and corrected supernatant (For G3 samples). Subsequentially, the islets were incubated for 30 minutes in KRBH buffer (500 μl per 10 islets) containing 20 mM glucose (For G20 samples). Subsequentially, the islets were incubated for 30 minutes in KRBH buffer (500 μl per 10 islets) containing 20 mM KCl (For KCl20 samples). At the end of the incubation period, the islets were pelleted by brief centrifugation, and an aliquot of the supernatant was collected. The secreted insulin in the supernatants was determined using the mouse insulin ELISA kit (Mercodia, 10-1247-01) for mouse islets, and the ultrasensitive human insulin ELISA kit (Mercodia, 10-1141-01) for human islets. For Insulin secretion assay from EndoC-βH1 cells, we split EndoC-βH1 cells 100,000 – 500,000 cells/well in Cultrex RGF Basement Membrane Extract-coated (4 μg/ml at 37°C for 1 hour pretreatment) 24 well plates. EndoC-βH1 cells were incubated in KRBH buffer (500 μl/well) supplemented with 3 mM glucose for 1 hour (G3) and then with 20 mM glucose (G20) for another 1 hour. Subsequently, the cells were incubated with 20 mM KCl/3 mM glucose (KCl20). The supernatant was collected after each stimulation. Intracellular and secreted insulin levels were determined by ultrasensitive Insulin ELISA Kit (Mercodia, 10-1132-01).

### Caspase 3/7 apoptosis assay

Caspase 3/7 cell apoptosis assays were performed using IncuCyte Caspase 3/7 green dye apoptosis reagent (Sartorius, 4440) and monitored using the IncuCyte SX5 live cell analysis instrument with according to the manufacturer’s protocol. Briefly, EndoC-βH1 cells or dispersed HILOs were plated overnight in 96-well plates (Thermo Scientific, 167008) at approximately 100,000 cells per well prior to the addition of small molecules and bile acids at the indicated concentrations. IncuCyte Caspase 3/7 green apoptosis reagent was added at a final concentration of 2.5 µM, with the cells being monitored for 240 hours. Live image analysis was performed using Sartorius IncuCyteSX5 live cell analysis software.

### Firefly luciferase assay

HEK293LTV cells or EndoC-βH1 cells were infected with an NFκB luciferase reporter lentivirus (pGreenFire1-NFκB; SBI, TR012PA-P). Approximately 100,000 cells per well were plated in 96 well-plates, pretreated with 10 μM FXR agonists or 500 nM to 1 μM BET inhibitors for 24 hours, and then treated with 10 ng/ml IL-1β for another 24 hours. The luciferase signal was measured using the Firefly Luciferase Assay Kit 2.0 (Biotium, 30085-2). Briefly, culture medium was aspirated from the plate and lysis buffer was added. Lysis was performed at 25°C for 15 minutes, before the addition of luciferase substrate. Luminescence was measured using a microplate reader (BioTek Synergy H1) within 10 minutes of substrate addition.

### siRNA transfection of EndoC-βH1

EndoC-βH1 with NF-κB luciferase reporter cell lines were seeded into 96 well plate (Corning Costar, 3610) at a density of 100,000 cells per well. Cells were then transfected with via lipofectamine 3000 transfection reagent (Invitrogen, L3000015), following manufacturer’s suggested protocol. The final concentration for siRNA for BRD2 (Santa Cruz Biotechnology cat#sc-60282), BRD3 (Santa Cruz Biotechnology, sc-60284), BRD4 (Santa Cruz Biotechnology, sc-43639), and control siRNA (Santa Cruz Biotechnology cat#sc-37007) were 100 nM. The cells were harvested after 24 hours to check for downregulation of target genes, or treated Fex (final concentration: 10 µM) or JQ1 (final concentration: 500 nM) for 24 hours, before IL-1β (final concentration: 10 ng/mL) was added. Cells were lysed and assayed for luciferase activity 24 hours post IL-1β treatment with Firefly Luciferase Assay Kit 2.0 (Biotium, 30085-2), following manufacturer’s suggested protocol.

### Gaussia luciferase assay for insulin secretion

EndoC-βH1 and INS-1 cell lines were infected with Proinsulin-NanoLuc lentivirus (Addgene, 62057). Approximately 100,000 cells were seeded in 96-well plates, pretreated with 10 μM FXR agonists or 500 nM to 1 μM BRD inhibitors for 24 hours, and then treated with 10 ng/ml IL-1β for another 24 hours. The insulin secretion assay was performed similar to the islet insulin secretion assay protocol described above. The supernatant was collected and assayed for luciferase signal using the Pierce Gaussia Luciferase Glow Assay Kit (Thermo Scientific, 16161) according to the manufacturer’s instructions. Luminescence was measured using a microplate reader (BioTek Synergy H1) within 10 minutes of substrate addition.

### RNA isolation and gene expression analysis

Cell lines were lysed using TRIzol reagent (Ambion, 15596018) and phenol-chloroform-isoamyl alcohol treatment, followed by total RNA extraction using the column of Aurum Total RNA Mini Kit (Bio-Rad, 732-6820) according to the manufacturer’s instructions. cDNA was synthesized from 500 - 1000 ng of total RNA treated with DNase I using the iScript cDNA Synthesis Kit (Bio-Rad, 1708891). Real-time quantitative RT-PCR (qPCR) was performed using the iTaq™ Universal SYBR Green Supermix (Bio-Rad, 1725122), and analyses were performed using the ABI7900HT Fast Real-Time PCR System or the ABI Step One Plus Real-Time PCR System. All samples were run in triplicate and mRNA levels were normalized to the universal reference gene, 36B4. Primers are listed in Supplementary Table.4.

### RNA-seq analyses

Total RNA was isolated from TRIzol (Ambion, 15596018) treated cell using the Aurum Total RNA Mini Kit (BIO-RAD, 7326820) and treated with DNase I for 30 minutes at room temperature (RT). Sequencing libraries preparation and sequencing was performed at Novogene, Inc. Short read sequences were mapped to a UCSC hg38 reference sequence using the RNA-Seq aligner STAR ^77^. The hg38 associated known splice junctions were provided to aligner, and the *de novo* junction discovery was also allowed. Differential gene expression analysis, statistical testing and annotation were performed using DESeq2. Genes with adjusted p value < 0.05 were considered significant. Additional fold-change thresholds were applied for visualization (|log2 fold change| > 0.5). Transcript expression was calculated as gene-level relative abundance in fragments per kilobase of exon model per million (fpkm) mapped fragments and correction for transcript abundance bias was applied ^78^. Gene ontology was performed using DAVID software. Heatmaps were generated using R-Script with heatmap.2 (gplot) software. The scale of heatmaps was determined by the Z-score. Motif analyses were performed using HOMER^79^

### ATAC-seq analyses

EndoC-βH1 cells were pretreated with DMSO, Fex (10 μM) for 72 hours and 10 ng/ml IL-1β for 48 hours. 100,000 cells were harvested to prepare ATAC-seq libraries. Nuclei were extracted from samples and tested for quality control (QC). After the QC, the nuclei pellets are resuspended in the Tn5 transposase reaction mix, which includes transposase and equimolar Adapter1 and Adapter 2. The mix of transposition reaction is incubated at 37°C for 30 min. PCR is then performed to amplify the library. Libraries were purified with the AMPure beads and library quality was assessed by Tapestation 4150. Sequencing was performed using Nova-Seq 6000 at Novogene. Reads were aligned by The Burrows-Wheeler Alignment algorithm (BWA) and peaks were called by MACS2 software. Homer was used for Motif analyses using default settings. Differential peaks were called by Homer using default settings (Fold change > 2, p value < 0.0001). Gene Ontology (GO) was performed using DAVID software.

### ChIP-seq analyses

EndoC-βH1 cells were pretreated with DMSO, Fex (10 μM) for 72 hours, and 10 ng/ml IL-1β for 48 hours. Next, 20 million cells were harvested for each ChIP assay. Cells were crosslinked with 2 mM disuccinimidyl glutarate (DSG) and fixed with 1% Paraformaldehyde (PFA). After fixation, nuclei from were isolated, lysed, and sheared with a Covaris Ultrasonicator ME220 to yield DNA fragment sizes of 200 to 1,000 bp followed by IP. Antibody used for ChIP-seq was BRD4 (Betyl laboratory, A301-985A50). Sequencing was performed using Illumina technology to generate paired-end reads for each sample. Sequence data quality was determined using FastQC software. Bowtie2 (v2.3.3.1) (–very-sensitive) was used to map ChIP-seq reads to the mouse reference genome GRCm38. Duplicate reads were filtered out using the MarkDuplicate function from Picard tools v.2.17.0 (http://broadinstitute.github.io/picard/). Reads per kilobase and million mapped read (RPKM)-normalized bigWig files were generated with bamCoverage from deepTools v3.3.2. ChIP-Seq peaks were called using findPeaks within HOMER using default parameters for TF (-style factor). De novo and known motif analyses were carried out using the findMotifsGenome.pl module of the Homer package with the “-size given” option.

### Co-immunoprecipitation and western blot

Cells were scraped and collected from the cell culture plate, followed by nuclei isolation with hypotonic solution (Cayman cat#10009301), and then lysed with ice-cold Pierce RIPA buffer (Thermo Scientific, 89900) containing Halt protease and phosphatase inhibitor cocktail (Thermo Scientific, 78442) for 15 minutes, and the centrifuged at 15,000 × g for 15 minutes. The supernatants were collected and incubated with 1 µg of anti-FXR or anti-BRD4 antibody for 4 hours at 4°C, and then precipitated with Dynabeads Protein G (Thermo Scientific, 10003D) for 1 hour at 4°C with rotation (15 rpm). Dynabeads were washed three times with ice-cold PBS, and the proteins were eluted by adding IgG elution buffer (Thermo Scientific, 21004), and LDS-sample buffer (Alfa Aesar, J61942.AD) for 5 minutes at 80°C. The eluted samples were separated on Bolt 4-12% Bis-Tris Plus Gel (Thermo Scientific, NW04125BOX) at 200 V for 22 minutes, and then transferred to an iBlot2 PVDF mini stack (Thermo Scientific, IB24002) using the iBlot2 transfer system. The membrane was blocked with 2% (w/v) nonfat dry milk (Apex, 20-241), washed five times with TBST (Bioland Scientific, TBST03-02) using SNAP i.d. 2.0 (Millipore) and then incubated in primary antibody overnight at 4°C. Clean-blot IP detection kit (Thermo Scientific, 21232) was used as secondary antibody to detect undenatured antibodies. The membrane was stained with ProSignal Femto (Prometheus, 20-302B) and visualized using the Bio-Rad ChemiDoc MP imaging system (Bio-Rad).

### Protein purification

20 – 30 million EndoC-βH1 cells or HEK293LTV cells were incubated with DMSO, 10 μμ Fex, 500 nM JQ1, or combination of Fex and JQ for 24 hours before being washed with PBS and collected in 50 mL tube. Afterwords, the nuclei were collected by incubation with 1 × hypotonic solution diluted from 10 × (Cayman Chemicals, #10009301) along with halt protease/phosphatase inhibitor cocktail (Thermo Scientific, #78442) for 30 minutes. NP100 alternative (millipoore cat#492016-100ML) was added to a final concentration of 1%, incubated for 15 minutes on ice, before being centrifuged at 14,000 × g for 30 seconds to collect the nuclei pellets. The nuclei pellets then were separated, and RIPA buffer containing protease/phosphatase inhibitor was added before sonication of the solution for 5 seconds. The resulting solution was centrifuged at 14,000 × g for 15 minutes followed by collection for the supernatant. Total protein was measured via Pierce BCA assay kit (Thermo Scientific, #23225). The total protein was normalized for each condition and treatment, and then incubated with 1 μg of antibody for 4 hours, before incubation with dynabeads protein G (Invitrogen, #10004D) for additional hour. The dynabeads were precipitated, collected and washed three times with ice cold PBS, transferred into a new tube, and stored in -80°C until processing for LC-MS/MS. Equivalent amounts of pull-down samples were treated with 8 M urea, 10 mM TCEP, 100 mM TEAB pH 8.5 to solubilize bound protein. Samples were diluted to 6 M urea using 100 mM TEAB, and digested overnight at 37°C with 0.04 μg of LysC (Promega - R-LysC), with constant rotation. Samples were subsequently diluted to 1 M urea with 100 mM TEAB, then incubated with 0.04 μg of Trypsin Gold (Promega) with continuous rotation. The resulting peptides were acidified to a final concentration of 2% formic acid (with 100% formic acid), isolated using a stacked C18/SCX STAGE tip, and eluted with 5% ammonium hydroxide, 80% acetonitrile. Elutions were evaporated to dryness in a Speedvac vacuum concentrator and redissolved in 0.1% FA in water. LC-MS/MS analyses were performed in UCI proteomic core.

### Structural visualization of FXR and BRD4

AlphaFold-predicted structures of human FXR (NR1H4; UniProt: Q96RI1) and BRD4 (UniProt: O60885) were obtained from the AlphaFold Protein Structure Database. The corresponding PDB files (NR1H4_AF-Q96RI1-F1-model_v4.pdb and BRD4_AF-O60885-F1-model_v4.pdb) were visualized using UCSF ChimeraX. Structural alignment and visualization were performed to examine the relative positioning of FXR lysine residues (K158 and K218) with respect to the bromodomain regions of BRD4. All structural interpretations are based on computational predictions and were used to support experimental findings.

### Statistics and Reproducibility

Statistical analyses were performed using Graphpad Prism (version 7-10) or Microsoft Excel. All quantitative data are presented as mean ± standard error of the mean (SEM), unless otherwise indicated. Statistical analyses were performed using GraphPad Prism version 10. Unpaired two-tailed t-tests, one-way ANOVA with Tukey’s multiple comparisons test, and two-way ANOVA were used as appropriate. Sample sizes, exact p values, and the number of biological replicates (*n*) are reported in the corresponding figure legends. For qPCR analyses, all data were obtained from biological triplicates, with each individual sample normalized using technical triplicates. All experiments were independently repeated at least three times with consistent results, unless otherwise stated. No data were excluded from the analyses, and no statistical methods were used to predetermine sample size. The investigators were not blinded to group allocation.

### Software and Programs for Bioinformatics Analysis

1. R studio
2. Venny 2.1(https://bioinfogp.cnb.csic.es/tools/venny/)
3. Functional annotation tool DAVID Bioinformatics Resources (https://david.ncifcrf.gov/summary.jsp)
4. HOMER^79^
5. GSEA^80^

## Acknowledgments

We thank Dr.Thomas Q de Aguiar Vallim for sharing FXR flox/flox mice. We thank the technical support from Meghan Tahbaz, Andrew Salib and Shudi Wang. This work was supported by grants from CTSI-UCLA awards UL1TR001881 (E.Y.), Thomas J. Beatson Jr. Foundation Grant #2022-006 (E.Y.), TRDRP research award T33IR6551 (E.Y.) and Allen Foundation, Inc 2024 (E.Y.). E.Y. is supported by JDRF Career Development Award (5-CDA-2022-1178-A-N). Research reported in this publication was supported by the National Institute of Diabetes and Digestive and Kidney Diseases of the National Institutes of Health under Award Number R01DK136888 (E.Y.). Z.W. is supported by R01 DK132651, UL1-TR002377, Roubos Family Fund in Research, and the Mayo NORP program. 1S10OD016328 from the National Institutes of Health (P.D.G.). J.C. and H.P. are supported by fellowship from the CIRM-training grant (EDUC4-12837). Y.H. is supported by Uehara foundation fellowship.

## Author Contributions

F.C., J.T., D.C., J.C., Y.H., N.P., C.T., J.W., L.W., Y.M., P.G., H.P., N.H., K.K., A.S., S.F., E.I., L.F.C., Z.W., and E.Y. performed experiments, analyze and interpreted results. F.C., J.W., Y.H., Z.W., and E.Y. performed bioinformatics and genome data analyses. P.G., L.F.C., Z.W., and E.Y. generated resources. F.C. and E.Y. drafted the original manuscript and all authors read and edited manuscript. E.Y. conceived and supervised study.

## Data and materials availability

RNA-Seq data supporting the results of this study have been deposited in the National Center for Biotechnology Information (NCBI) Sequence Read Archive (SRA) database, accession BioProject ID: PRJNA1046214.

---

**Supplementary Fig. 1.**
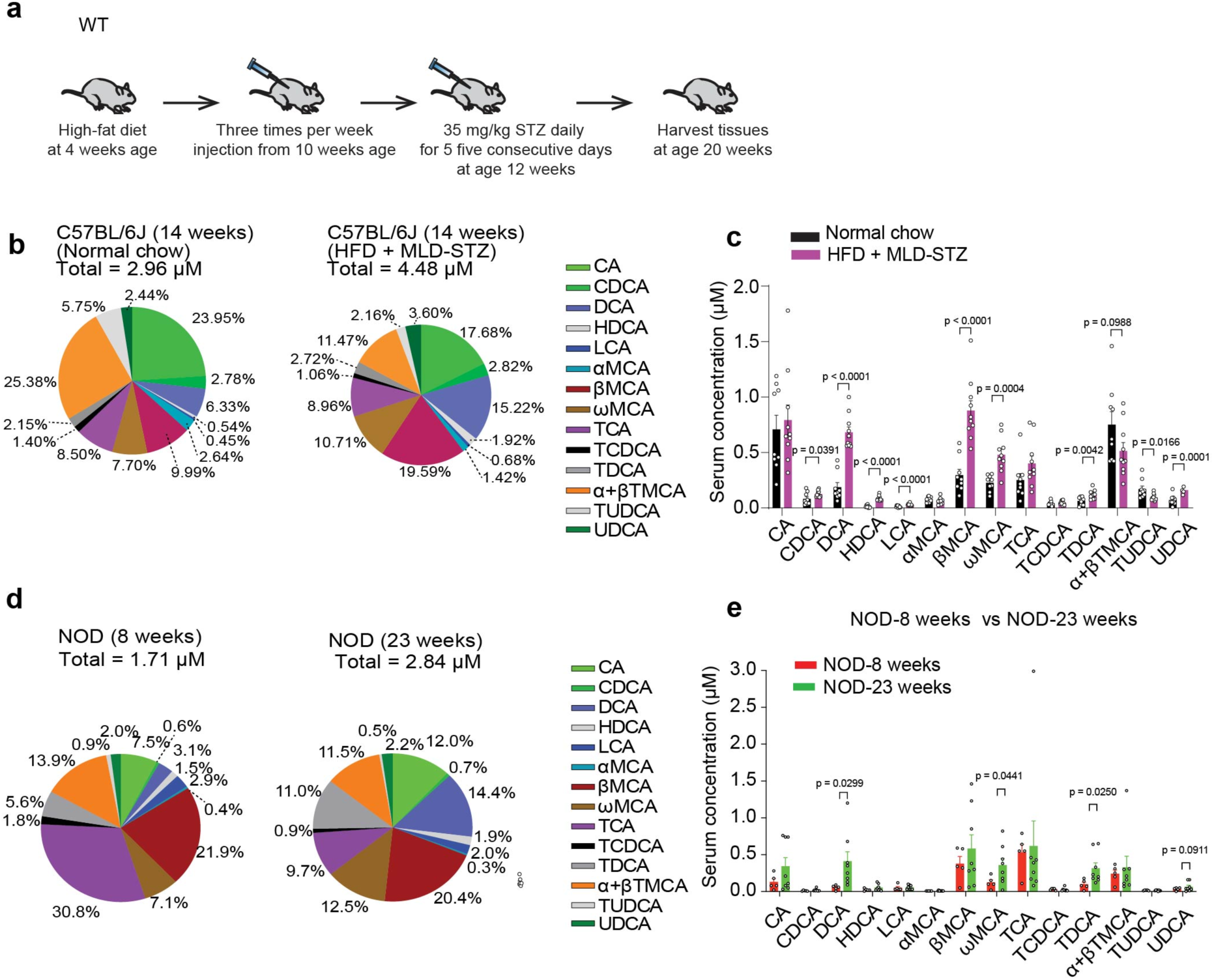
Dysregulation of bile acid signaling in HFD + MLD-STZ and NOD mouse models. **a,** Schematic of the T2D model. C57BL/6J mice were fed a high-fat diet (HFD) starting at 4 weeks of age. Pre-treatment was initiated at 10 weeks, followed by multiple low-dose STZ (MLD-STZ) administration for five consecutive days starting at 12 weeks. **b**, Comparison of serum bile acid (BA) profiles shown as total levels and relative composition between non-diabetic, age-matched male C57BL/6J mice fed a normal chow diet (left pie chart) and diabetic, age-matched male C57BL/6J mice subjected to HFD + MLD-STZ treatment (right pie chart). Serum samples were collected 14 days after the final STZ injection. n = 9 (normal chow), n = 10 (HFD + MLD-STZ).**c**, Comparison of serum individual BA amount in non-diabetic, normal chow-fed C57BL/6J versus diabetic C57BL/6J on HFD + MLD-STZ. n = 9 for C57BL/6J on normal chow, n = 10 for C57BL/6J on HFD + MLD-STZ. **d**, Serum bile acid (BA) profile of NOD mice (8 weeks or 23 weeks, female) measured by HPLC-MS/MS. e, Comparison of serum BA amounts in female NOD mice at 8 weeks vs 23 weeks. 8 weeks NOD n = 5, 23 weeks NOD n = 8. Statistical analysis was performed using unpaired two-tailed Student’s t-test. Error bars represent mean ± SEM.

**Supplementary Fig. 2.**
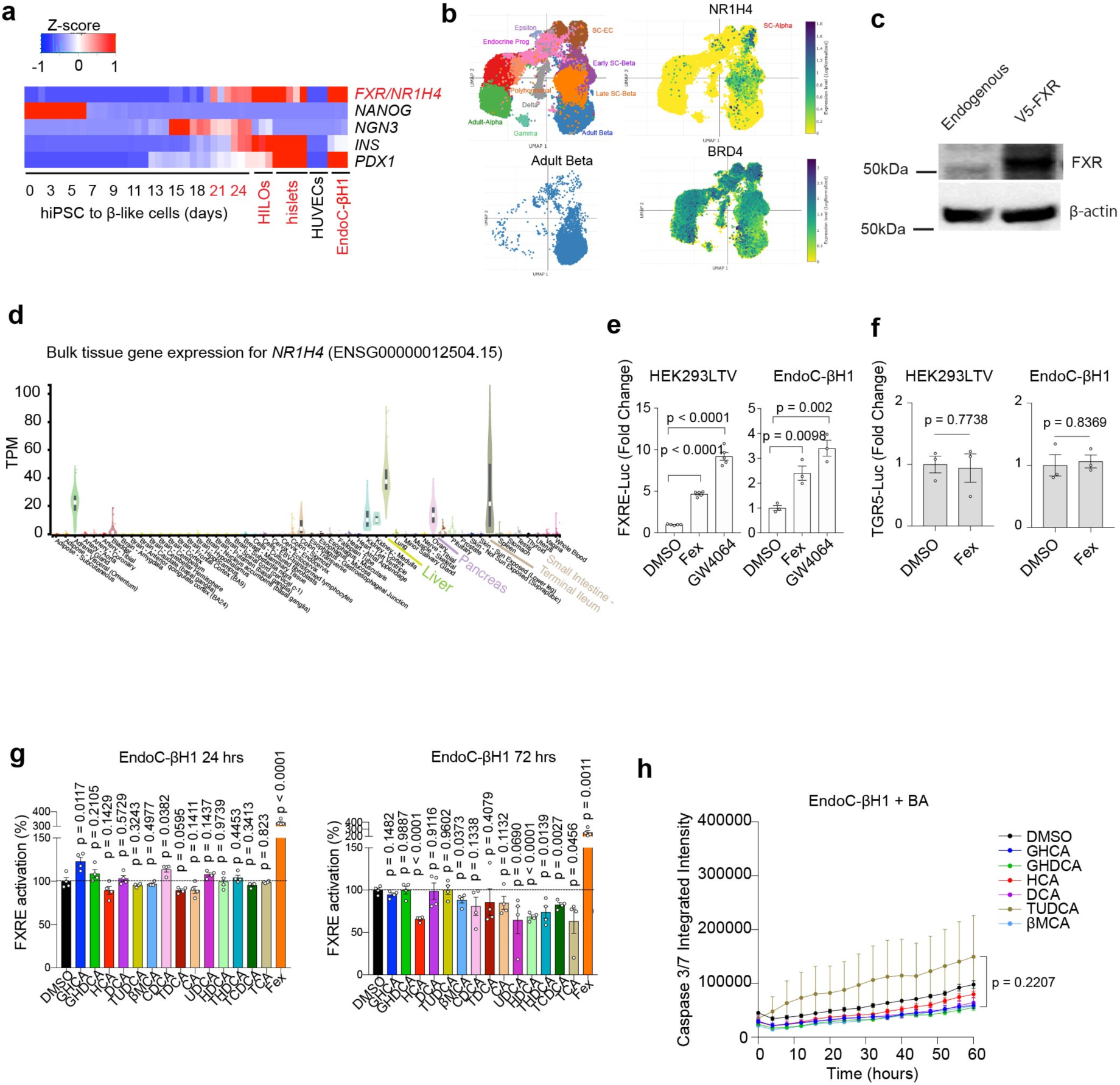
FXR signaling in human β cells. **a**, Heatmap of FXR gene expression and β cell marker genes in human iPSCs during their stepwise differentiation into HILOs compared to human islets, HUVEC cells, and EndoC-βH1. **b**, *NR1H4* gene expression in human islets and hPSC-derived pancreatic cells. Data was obtained from single cell Portal Broad institute (Functional, metabolic and transcriptional maturation of human pancreatic islets derived from stem cells). **c**, Representative image of FXR Western blot for endogenous and DOX-induced FXR overexpression system in EndoC-βH1 human β cell line. **d**, *NR1H4* expression in human tissues. Data was obtained from GTEx Portal. **e**, FXR reporter assay after 24 hours of treatment with DMSO, Fex (10 μM) or GW4064 (10 μM) in HEK293LTV and EndoC-βH1 cells. n = 3/each. **f**, TGR5-luciferase in EndoC-βH1 and HEK293LTV cell lines containing the TGR5 luciferase reporter after 24 hours treatment with the FXR-specific agonist Fex. n = 3/each. **g**, FXRE (FXR responsive element)-luciferase reporter signal of different BAs (10 μM each for indicated time) and Fex on EndoC-βH1 cells after 24 hours and 72 hours treatment. n = 4/each. **h**, Real-time imaging of caspase 3/7 green intensity signal in EndoC-βH1 cell line for 60 hours incubation with BAs (10 μM each for indicated time) without inflammatory cytokine IL-1β. n = 4/each. Statistical analyses were performed using unpaired Student’s two-tailed t-tests for (f), one-way ANOVA with Tukey’s multiple-comparison test for (e, g), and two-way ANOVA for (h). Error bars represent mean ± SEM.

**Supplementary Fig. 3.**
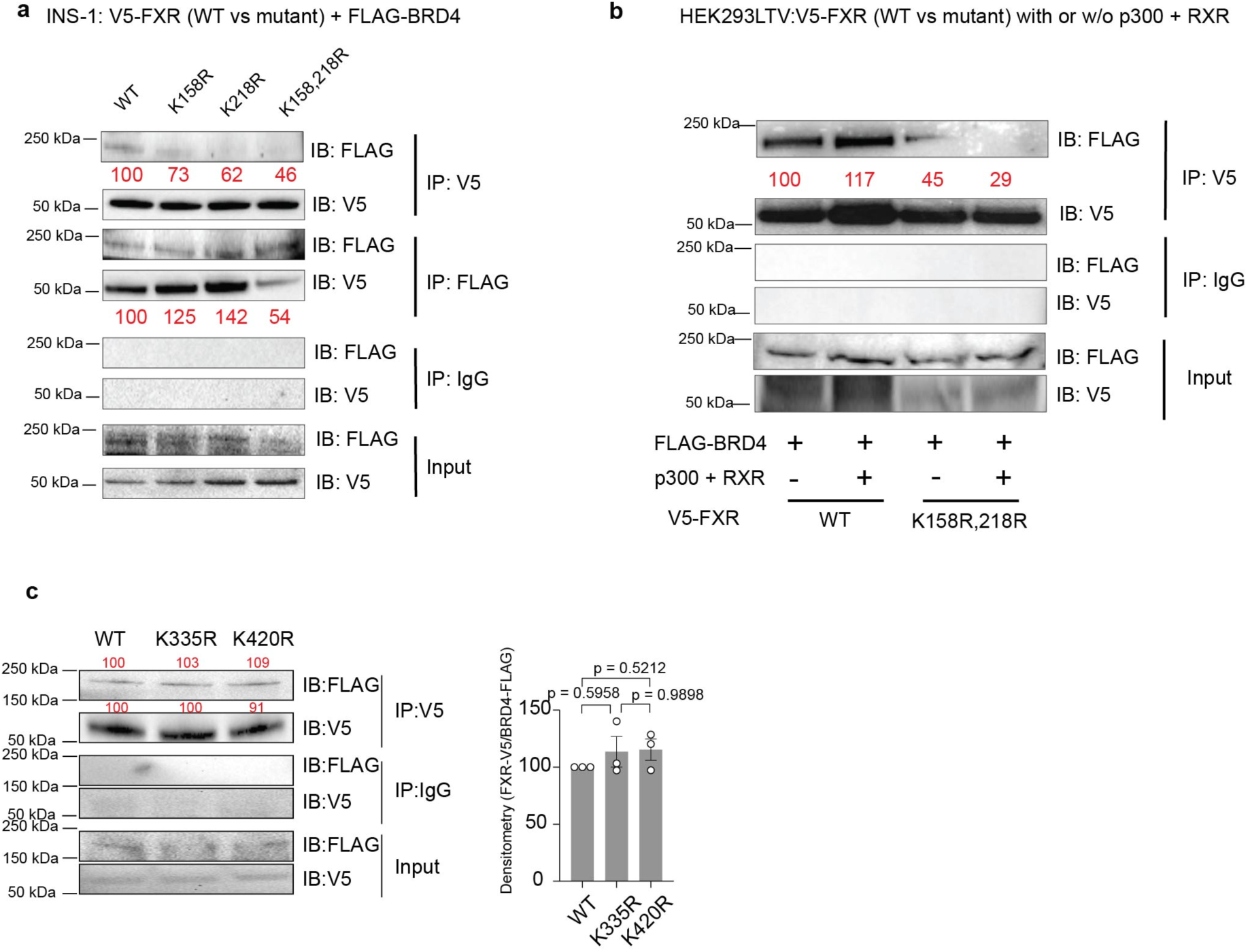
Physical interaction between FXR and BRD2, BRD3, or BRD4. **a**, IB: FLAG-BRD4 or V5 in V5-FXR (control), V5-FXR (K158R), V5-FXR (K218R), V5-FXR (K158R/K218R) pull-down (IP:V5, FLAG or IgG) protein complexes from INS-1 cells. **b**, Co-IP of FLAG-BRD4 or V5-FXR + p300 + RXR in HEK293LTV cells after co-transfected with FLAG-BRD4 and V5-FXR (WT) or V5-FXR (mut, K158R/K218R). **c**, Immunoblot (IB) of FLAG or V5 in V5-FXR (WT), V5-FXR (K335R), V5-FXR (K420R) pull-down (IP:V5) protein complexes from EndoC-βH1 cells. Red numbers above the blots indicate densitometry values presented as percent relative to FXR-WT which is set as 100. The bar graph represents densitometric analyses of FXR-V5/BRD4-FLAG ratios from three independent experiments. Statistical analysis was performed using one-way ANOVA with Tukey’s multiple-comparison test. Error bars represent mean ± SEM

**Supplementary Fig. 4.**
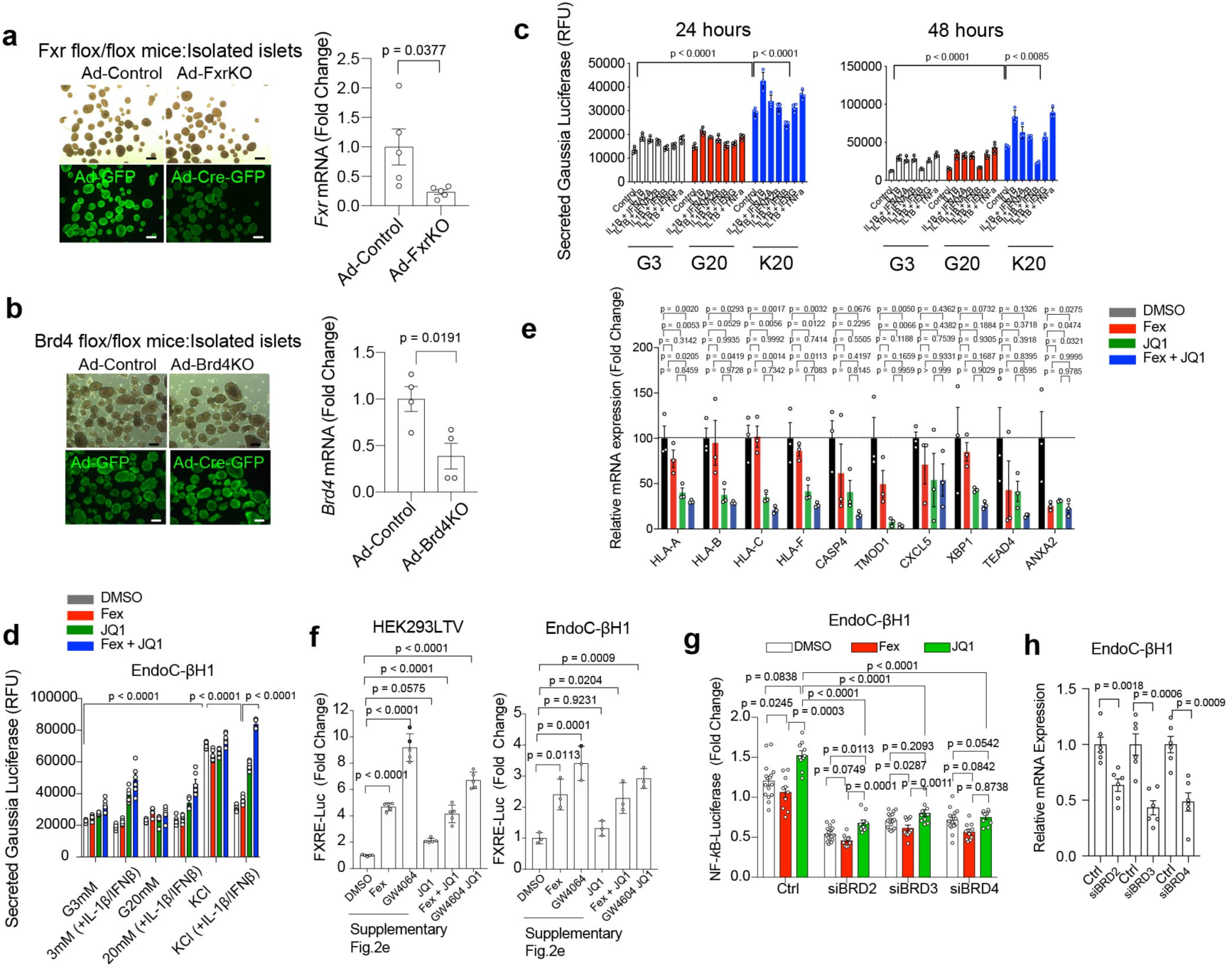
FXR agonist and BET inhibitor synergistically regulate β cell insulin secretion via FXR and BRD4 under inflammatory stress. **a**, Upper panel: Optical and fluorescence image of isolated islets from Fxr flox/flox mice with adenovirus-mediated GFP (Ad-Control) or Cre-GFP (Ad-FxrKO) expression. Bottom panel: qPCR analyses of *Fxr* gene expression in Ad-Control and Ad-FxrKO islets. Scale bar = 100 µm. **b**, Upper panel: Optical and fluorescence image of isolated islets from Brd4 flox/flox mice with adenovirus-mediated GFP (Ad-Control) or Cre-GFP (Ad-Brd4KO) expression. Lower panel: qPCR analyses of *Fxr* gene expression in Ad-Control and Ad-Brd4KO islets. Scale bar = 100 µm. **c**, Gaussia-luciferase reporter assay for insulin secretion of EndoC-βH1 cells after 48 hours incubation with DMSO, Fex (10 μM), JQ1 (500 nM) or Fex (10 μM) + JQ1 (500 nM) and with or without 24 hours or 48 hours treatment with the inflammatory cytokines IL-1β (10 ng/ml) and IFNα2A (10 ng/ml), IFNα2B (10 ng/ml), IFNβ (10 ng/ml), IFNγ (10 ng/ml) or TNFα (10 ng/ml). n = 6/each. **d**, qPCR analyses of indicated genes in EndoC-βH1 cells pretreated with DMSO, Fex (10 μM), and/or JQ1 (500 nM) for 72 hours and with IL-1β (10 ng/ml) + IFNγ (10 ng/ml) for 48 hours. n = 3/each. **e**, Gaussia-luciferase reporter assay for insulin secretion corresponding to Fig. 3e. **f**, FXR reporter assay of EndoC-βH1 and HEK293LTV cells after 24 hours of treatment with Fex (10 μM), GW4604 (10 μM) and/or JQ1 (500 nM). n=3/each. **g**, NF-*k*B luciferase activity in EndoC-βH1 cells transfected siRNA against *BRD2* (siBRD2), *BRD3* (siBRD3), or *BRD4* (siBRD4) respectively 96 hours prior of the Luciferase assay. Fex (10 μμ) or JQ1 (500 nM) were treated for 48 hours and IL-1β (10 ng/ml) for 24 hours prior of the Luciferase assay. **h**, qPCR analyses of BRD2, BRD3, or BRD4 48 hours after siRNA transfection in EndoC-βH1 cells. Statistical analyses were performed using unpaired two-tailed Student’s t-tests for (a, b, h) and one-way ANOVA with Tukey’s multiple-comparison test for (c–g). Error bars represent mean ± SEM.

**Supplementary Fig. 5.**
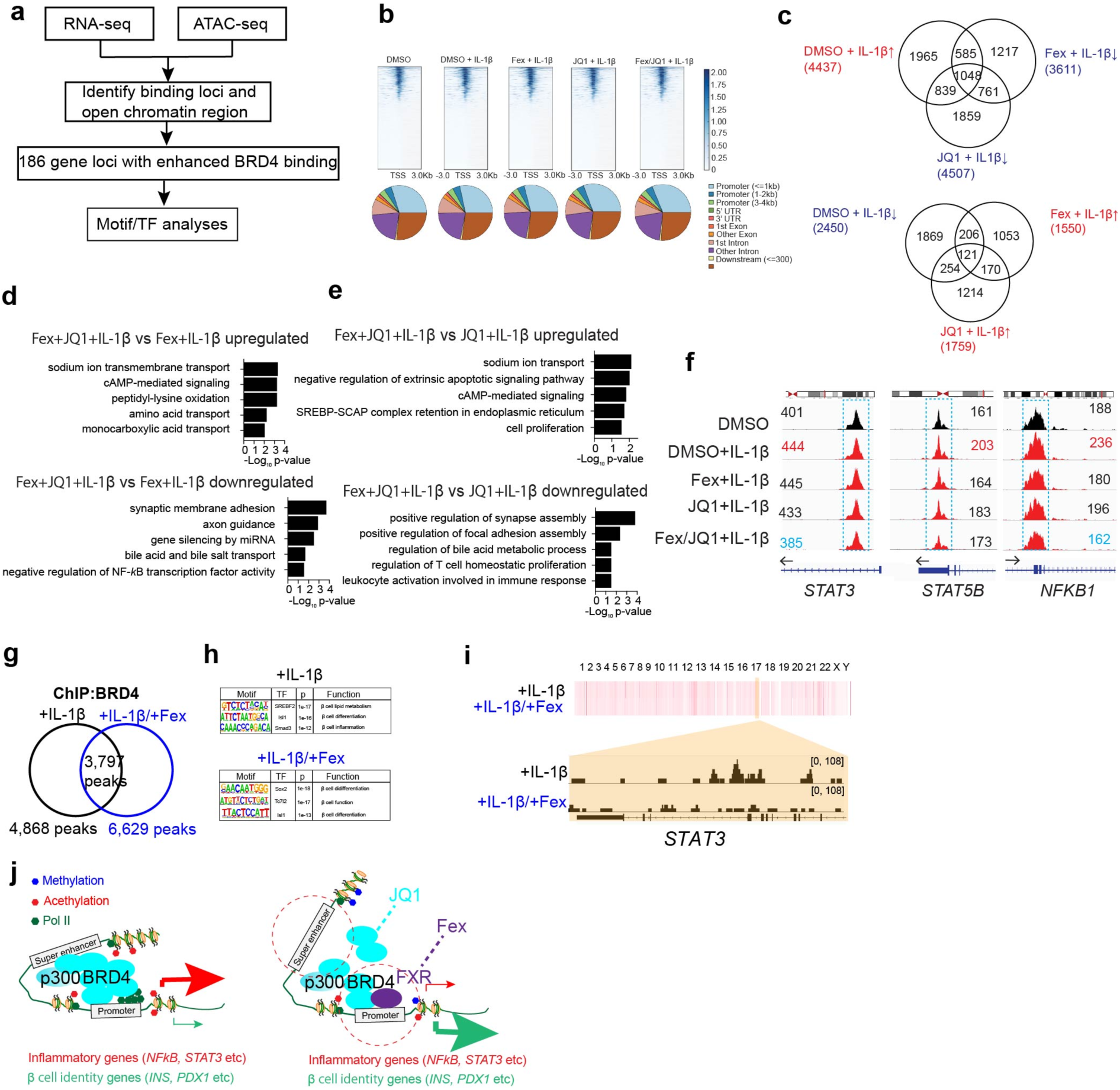
ATAC-seq and BRD4 ChIP-seq analyses in human β cells. **a**, Schematic of RNA-seq, ATAC-seq and ChIP-seq of BRD4 binding loci identification in EndoC-βH1 cells. **b**, Chromatin accessibility at sites in response to DMSO, DMSO + IL-1β, Fex + IL-1β, JQ1 + IL-1β, or Fex + JQ1 + IL-1β in EndoC-βH1 cells. **c**, Number of peaks significantly up or downregulated by IL-1β, while preserved by Fex or JQ1. **d, e**, GO of synergistically up or down regulated by Fex + JQ1 compared to single treatment of Fex (**d**) or JQ1 (**e**) under the stress of IL-1β. **f**, Browser tracks showing ATAC-seq data with indicated treatments at indicated loci. **g**, Venn diagram for BRD4 binding regions at IL-1β (10 ng/ml, 24 hours) with or without Fex (10 μM, 48 hours) treatment in EndoC-βH1 cells. **h**, Motif analysis of BRD4 binding regions in EndoC-βH1 cells. **i**, Genome browser view of BRD4 binding to the *STAT3* locus in IL-1β (10 ng/ml, 24 hours) with or without Fex (10 μM, 48 hours) treated EndoC-βH1 cells. **j**, Model of BRD4 and FXR mediated transcriptional regulation for inflammatory and β cell identity genes.

**Supplementary Fig. 6.**
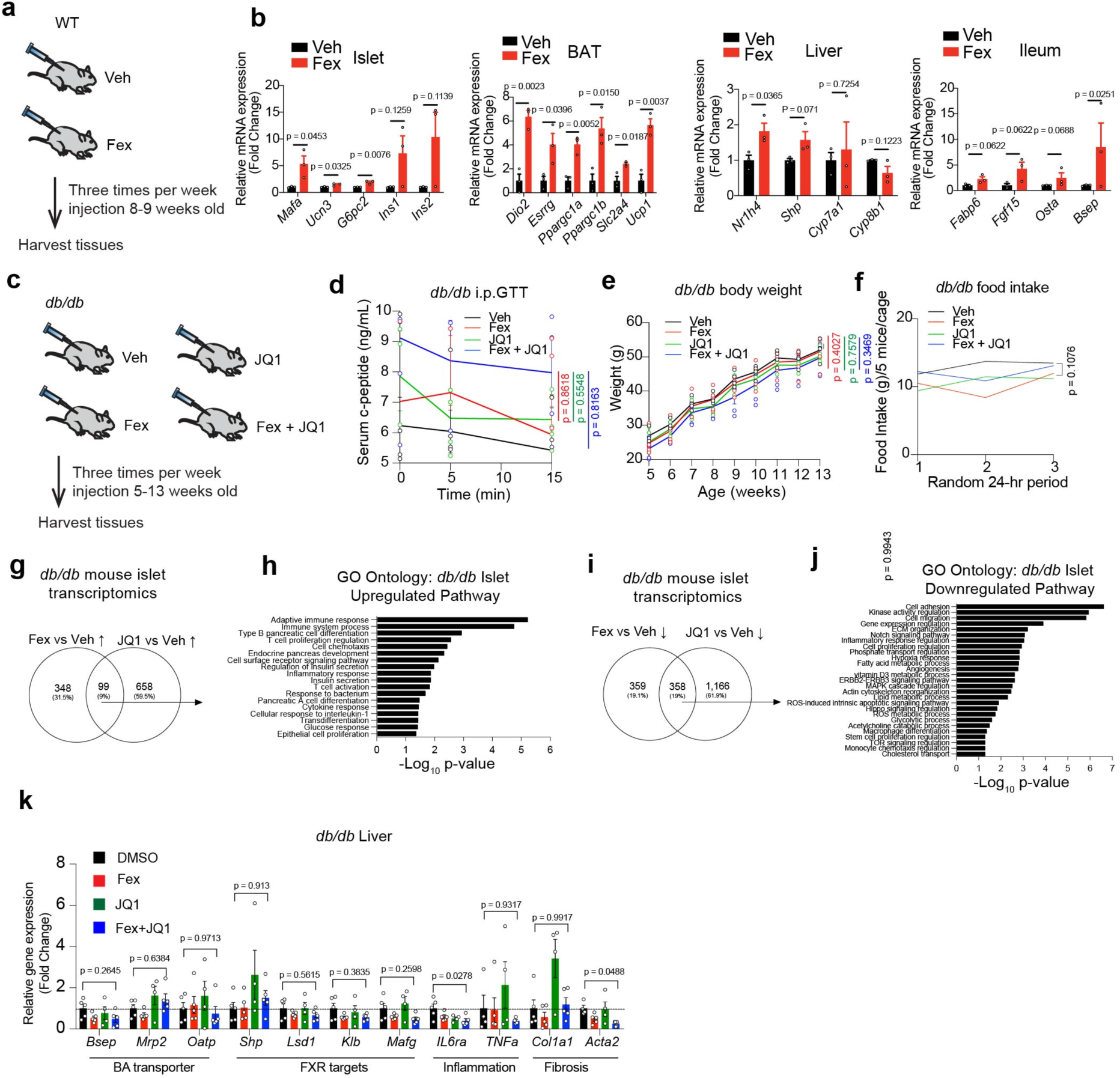
FXR agonist and BET inhibitor do not alter body weight gain in *db/db* mice. **a,** Scheme of mouse experiments. Veh or Fex (50 mg/kg) are i.p. injected three times a week between 8 to 9 weeks old C57BL6J male mice and tissue are harvested 24 hours after the last i.p.. **b,** qPCR analyses of indicated genes in isolated islets, brown adipose tissue (BAT), liver and ileum. n = 5 each. **c,** Scheme of mouse experiments. Veh, Fex (50 mg/kg), JQ1 (10 mg/kg), or Fex (50 mg/kg) + JQ1 (10 mg/kg) are i.p. injected daily from 5 weeks to 13 weeks old *db/db* male mice and tissue are harvested. **d,** Glucose tolerance test for serum c-peptide of *db/db* after 8-week i.p. treatment with Veh, Fex, JQ1 or Fex+JQ1. n = 5/each. **e**, Body weight of *db/db* mice during the 8-week i.p. treatment with Veh, Fex, JQ1 or Fex+JQ1. n=5/each. **f**, Food consumption of *db/db* mice at three random 24 hour periods during the 8-week i.p. treatment with Veh, Fex, JQ1, or Fex+JQ1. n = 5/each. **g**, Venn diagram of differentially upregulated genes in pancreatic islets isolated from *db/db* mice treated with Fex or JQ1. n=3/each. **h**, Gene ontology pathway analysis of the common 99 differentially upregulated genes in pancreatic islets isolated from *db/db* mice treated with Fex or JQ1. n = 3/each. **i**, Venn diagram of differentially downregulated genes in pancreatic islets isolated from *db/db* mice treated with Fex or JQ1. n = 3/each. **j**, Gene ontology pathway analysis of the common 358 differentially downregulated genes in pancreatic islets isolated from *db/db* mice treated with Fex or JQ1. n = 3/each. **k**, qPCR analyses of indicated genes in liver of *db/db* at 13 weeks of age. Statistical analyses were performed using unpaired two-tailed Student’s t-tests for (b) one-way ANOVA with Tukey’s multiple-comparison test for (f, k) and two-way ANOVA for (d, e). Error bars represent mean ± SEM.

**Supplementary Fig. 7.**
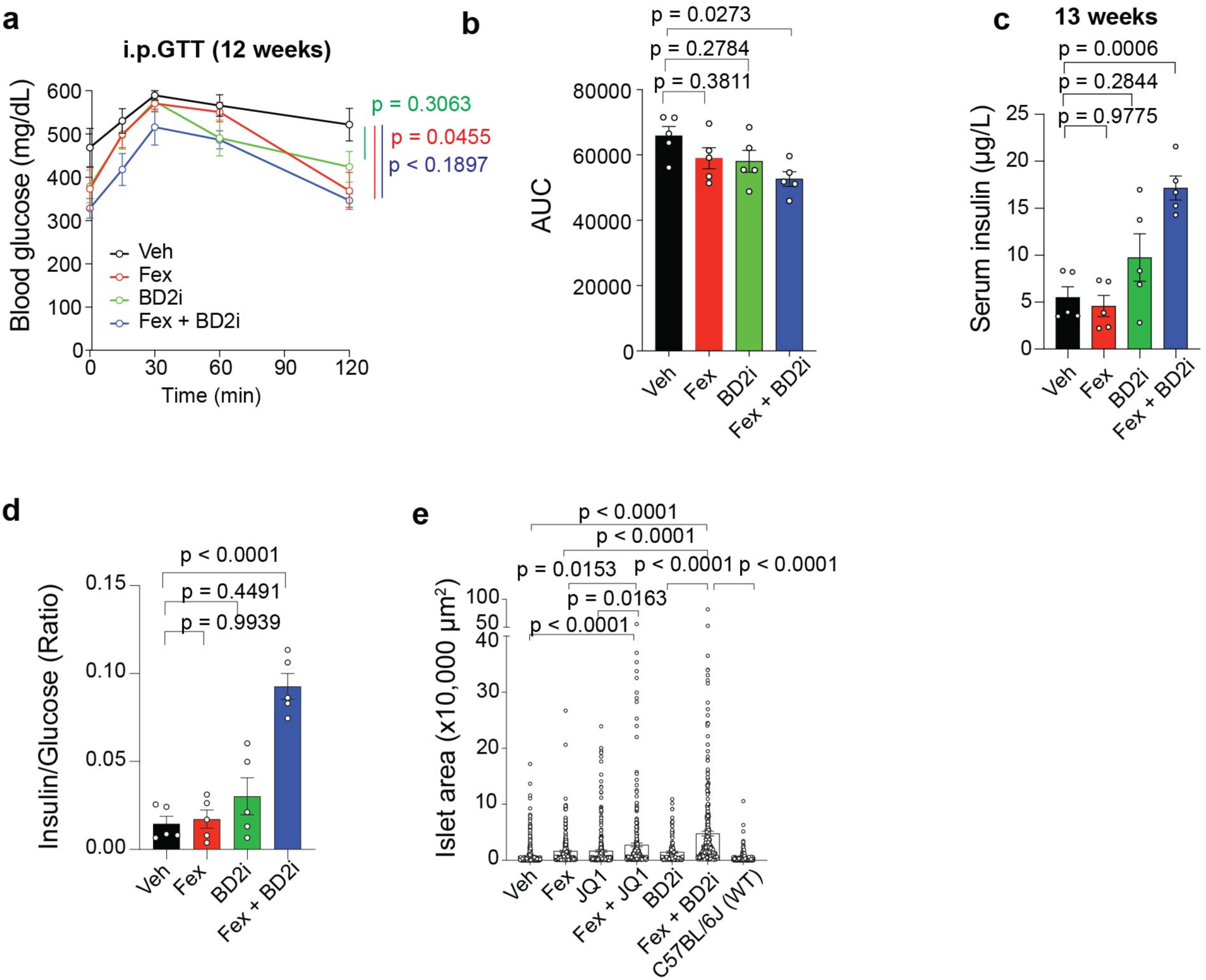
FXR agonist and iBET-BD2 inhibitor improve glucose homeostasis in *db/db* mice. **a**, Intraperitoneal glucose tolerance test on the 12-week old of i.p. treatment with Veh, Fex, JQ1 or Fex + JQ1 combination. n = 5/each. **b**, Area under the curve (AUC) for the intraperitoneal glucose tolerance test at the 8-week of i.p. treatment with Veh, Fex, JQ1 or Fex + JQ1 combination. n = 5/each. **c**, Serum insulin/glucose index of 12 week old *db/db* with i.p. treatment with Veh, Fex, JQ1 or Fex + JQ1 combination. n = 5/each. **d**, Serum insulin/glucose index of *db/db* mice i.p. treatment of Veh, Fex, JQ1 or Fex + JQ1 combination. n = 5/each. **e**, Area of islets from H&E staining of *db/db* mouse pancreas after 8-weeks of i.p. treatment with vehicle, Fex, JQ1, BD2i, Fex + JQ1 or Fex + BD2i combination. Two histology slides per condition. Statistical analyses were performed using two-way ANOVA for (a) and one-way ANOVA with Tukey’s multiple-comparison test for (b–e). Error bars represent ± SEM.

**Supplementary Fig. 8.**
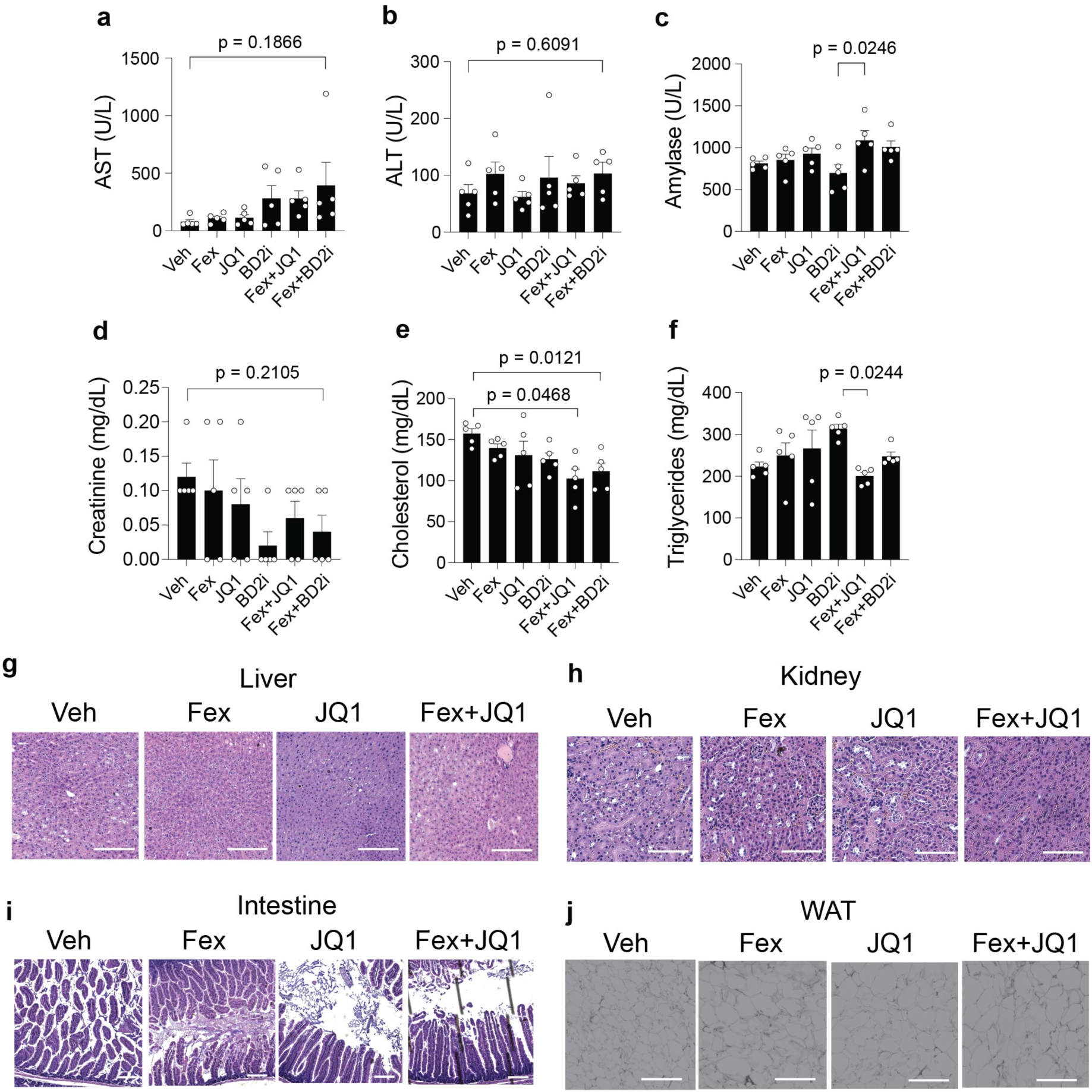
Impact of serum parameters by FXR agonist and BET inhibitors in *db/db* mice. (**a-f**), Serum metabolic parameters are measured at 13 weeks old *db/db* mice after treating with indicated drugs with daily i.p. injection from 5 weeks old to 13 weeks old. AST (U/L) (**a**), ALT (U/L) (**b**), Amylase (U/L) (**c**), Creatinine (mg/dL) (**d**), Cholesterol (mg/dL) (**e**) and Triglycerides (**f**). (**g-j**) Representative images for H&E stain for the liver (g), kidney (h), intestine (i) and white adipose tissue (WAT) (j) of *db/db* mice after the 8-week of i.p. treatment with Veh, Fex, JQ1 or Fex + JQ1 combination. scale bar = 100 µm. Statistical analyses were performed using one-way ANOVA with Tukey’s multiple-comparison test (a-f). Error bars represent mean ± SEM.

**Supplementary Fig. 9.**
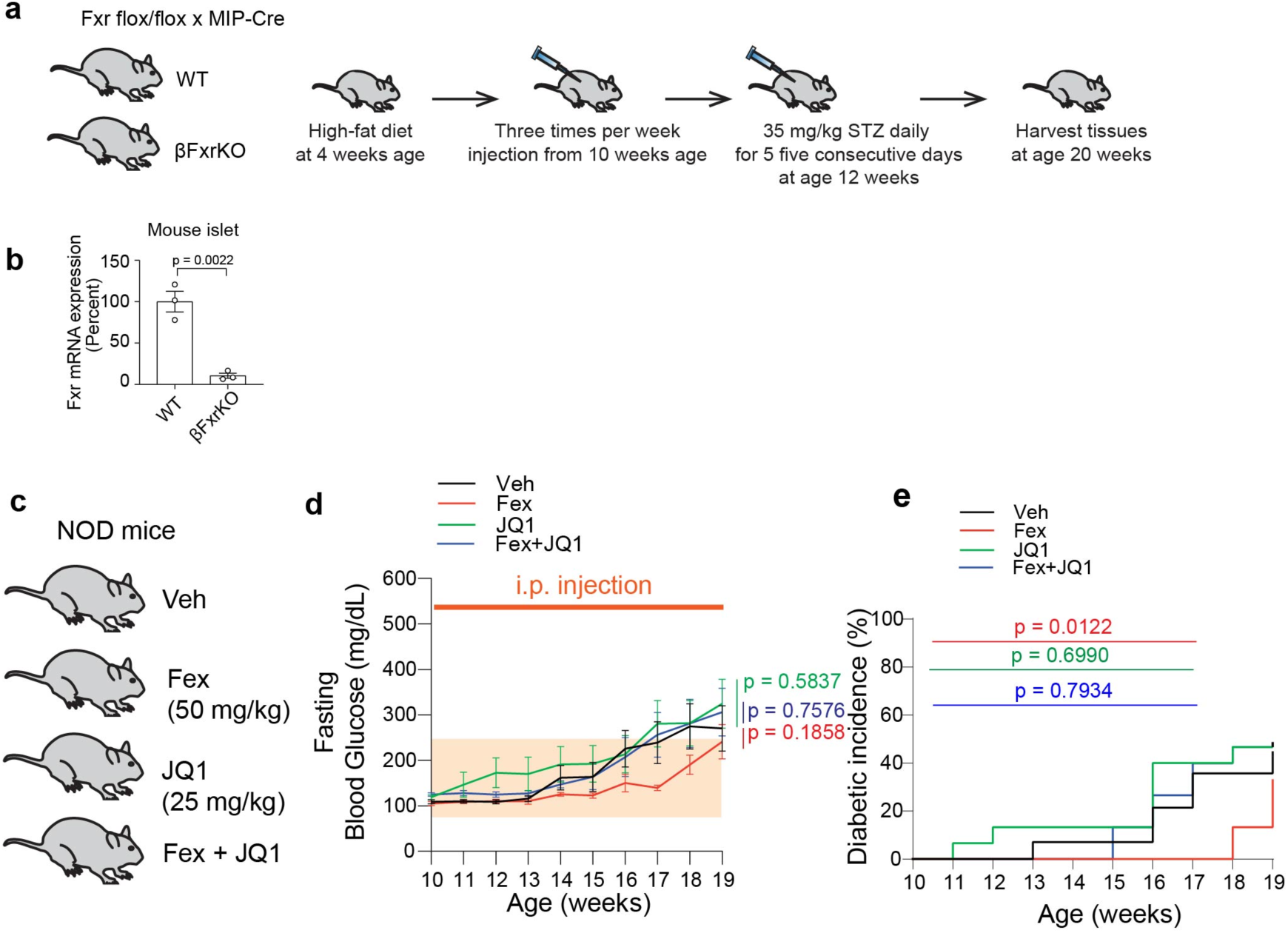
FXR agonist slow-down the progression of spontaneous T1D in NOD mice. **a,** Schematic of the T2D model. C57BL/6J or βFxrKO mice were fed a HFD starting at 4 weeks of age. Pre-treatment was initiated at 10 weeks, followed by MLD-STZ administration for five consecutive days starting at 12 weeks. **b**, qPCR analysis of Fxr target gene expression in isolated islets from WT and βFxrKO mice. **c**, Scheme of mouse experiments. Veh, Fex (50 mg/kg), JQ1 (25 mg/kg) or Fex (50 mg/kg) + JQ1 (25 mg/kg) are i.p. injected daily from 10 weeks to 19 weeks old NOD female mice. **d**, Fed *ad lib* blood glucose levels (mg/dL). n = 5. **e**, Diabetic incidents (% > 250 mg/dL blood glucose levels). Statistical analyses were performed using unpaired two-tailed Student’s t-test for (b), two-way ANOVA for (d), and Kaplan–Meier analysis with the log-rank (Mantel–Cox) test for (e). Error bars represent ± SEM.

**Supplementary Fig. 10.**
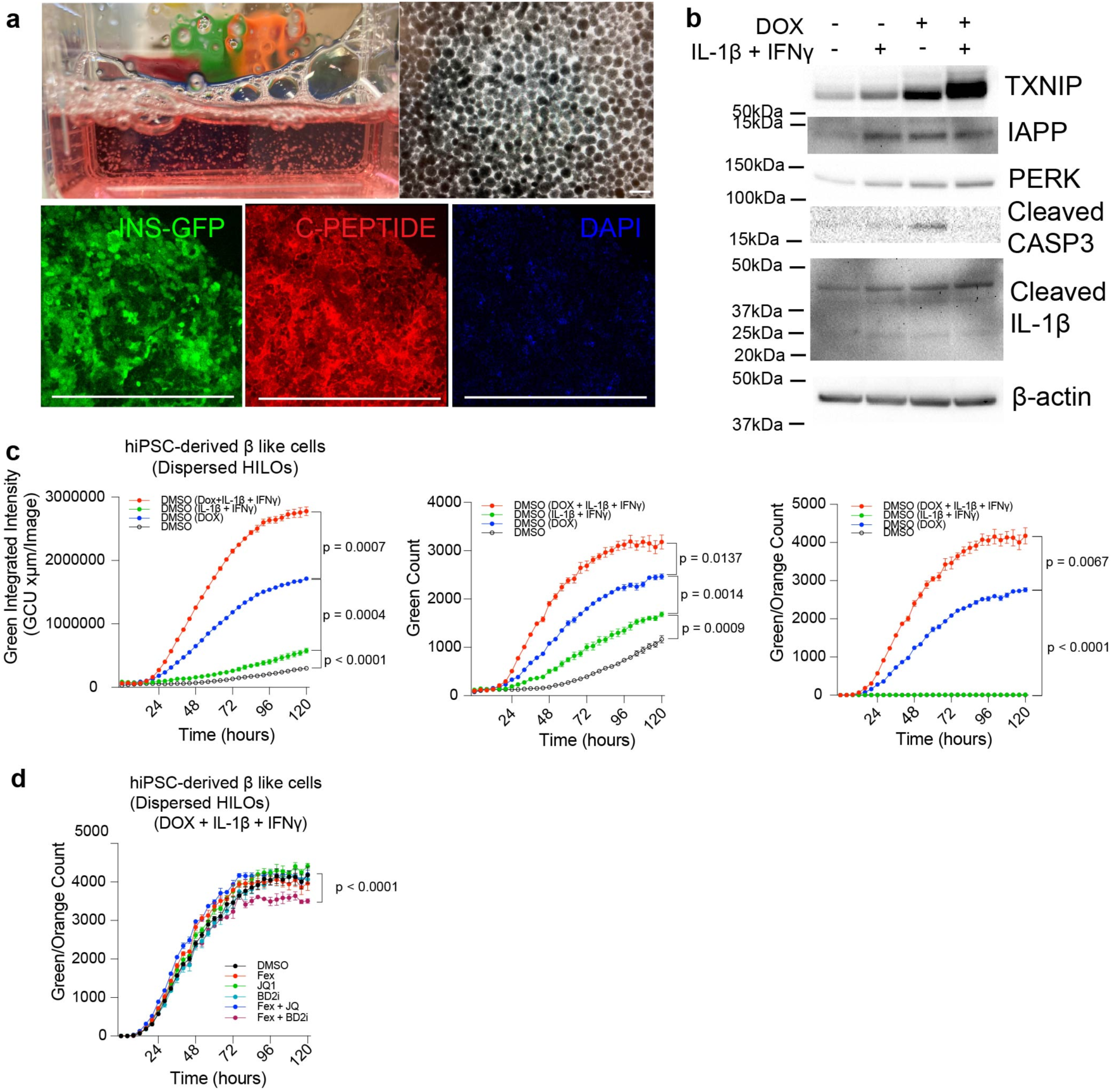
Model of human T2D in HILOs. **a**, Representative image of 3D culture system to generate HILOs. Scale bar = 50 µm. **b**, DOX-inducible human TXNIP and human IAPP overexpression in dispersed HILOs. IB for TXNIP, Amylin (IAPP), PERK, cleaved caspase 3 and β-actin. **c**, Apoptosis rate in dispersed HILOs was measured by real-time imaging using Incucyte SX5. Dispersed HILOs were treated with DOX (1 µg/µl), IL-1β (10 ng/ml), and IFNγ (10 ng/ml) for 120 hours. n = 3/each. Apoptosis rate was determined by green-integrated intensity, green count and orange-green count (DOX-inducible TXNIP/IAPP coexpress mCherry and it’s overlapping to Caspase 3/7 green fluorescence) respectively. n = 3/each. **d**, Apoptosis rate in dispersed HILOs was measured by real-time imaging using Incucyte SX5. Dispersed HILOs were treated with Fex (10 µM), JQ1 (500 nM), or iBET-BD2 (1 µM) under the stimulation of DOX (1 µg/µl), IL-1β (10 ng/ml), and IFNγ (10 ng/ml) for 120 hours. n = 3/each. Apoptosis rate was determined by orange-green count (DOX-inducible TXNIP/IAPP coexpress mCherry, which is colocalized with Caspase 3/7 green fluorescence) respectively. n = 3/each. Statistical analyses were performed using one-way ANOVA with Tukey’s multiple-comparison test for (c, d). Error bars represent mean ± SEM.

